# Structural Reorganization Following a Brain Tumor: A Machine Learning Study Considering Desynchronized Functional Oscillations

**DOI:** 10.1101/2022.11.14.516248

**Authors:** Joan Falcó-Roget, Fabio Sambataro, Alberto Cacciola, Alessandro Crimi

## Abstract

Neuroimaging studies have allowed for non-invasive mapping of brain networks in brain tumors. Although tumor core and oedema are easily identifiable using standard MRI acquisitions, imaging studies often neglect signals, structures and functions within their presence. Therefore, both functional and diffusion signals, as well as their relationship with global patterns of connectivity reorganization, are poorly understood. Here, we explore functional activity and the structure of white matter fibers considering the contribution of the whole tumor in a surgical context. First, we find that intra-tumor signals do exist and are correlated with alterations present both in healthy tissue and resting-state networks. Second, we propose a fiber tracking pipeline capable of using anatomical information while still reconstructing bundles in tumoral and peritumoral tissue. Finally, using machine learning and healthy anatomical information, we predict structural rearrangement after surgery given the preoperative brain network. The generative model also disentangles complex patterns of connectivity reorganization for different types of tumors. Overall, we show the importance of carefully designing studies including MR signals within damaged brain tissues, as they exhibit and relate to non-trivial patterns of both structural and functional (dis-)connections or activity.

## Introduction

Functional networks of brain tumor patients, as obtained from functional magnetic resonance imaging (fMRI), show clear altered patterns of local and global disconnections [1] that follow functional rather than spatial distance to tumor location [2]. Consistent with these findings, a separate study also reported functional abnormalities that overlap with unaffected structural areas [3]. Large-scale theoretical models of functional activity also report differences in inhibitory connections between networks having similar structural features [4, 5]. However, all these findings do not address the question of how the functional signal itself is modified by the presence of the tumor nor how it relates to potential desynchronizations in resting-state networks.

The oscillatory nature of functional time series has been widely exploited to entangle brain regions involved in a wide range of cognitive scenarios [6]. These functional co-activations, or connections [7], are computed by assessing temporal correlations which, by mathematical construction, rely on the oscillatory frequencies present in the signals. We hypothesized that the level of participation of each of those frequencies would be severely modified by the presence of a brain tumor. If this were true, we could quantify how these deviations propagate, or at least correlate, to alterations in the functional connections of resting-state networks similarly to what was shown for a system of coupled oscillators [8, 9]. Thereupon, we analyze local and brain-wide Fourier transformed functional time series and study how their power spectrum singularities account for network anomalies in brain tumor patients.

A parallel line of research has recently focused on the use of diffusion magnetic resonance imaging (dMRI) in a wide variety of brain diseases [10, 11]. Despite promising progress in the use of dMRI to resolve brain tumor microstructure [12], tumoral tissue is usually contaminated with cerebrospinal fluid and gray matter abnormalities, therefore posing a challenge to existing fiber reconstruction methods. Although several intra-lesion fiber tracking methods have been developed, their usage is still scarce. Successfully removing the contribution of cerebrospinal fluid requires the use of low angular resolution tensor diffusion models unable to resolve complex white matter regions [13, 14], or disregarding multi-shell acquisition schemes which are able to improve fiber orientation estimation when employing for instance constrained spherical deconvolution approaches [15]. Arguably, the lack of acceptance may be built upon the detrimental effects of disregarding the aforementioned MRI acquisition protocols and/or diffusion signal modeling approaches [16], due to longer acquisition time that is often not suitable in the clinical setting. We propose and describe a hybrid pipeline using a single-shell-3-tissue algorithm [17] only in the tumoral area of the brain. Afterwards, we merge the tractogram with the one obtained using multi-shell algorithms in healthy tissue, exploiting the best features of each method and minimizing the impact of their downsides.

Unlike functional network studies, the knowledge of structural networks in brain tumors is, to some extent, unclear. Previous groups failed to find significant tumor-dependent differences between patients and healthy subjects [18, 4], suggesting that network integration and segregation are mostly preserved. However, gliomas significantly altered the structural topology of the ipsilesional but no the contralesional hemisphere [19]. In addition, small structural differences vanished after surgery [5], implying that high-precision surgical interventions allow for spontaneous recovery of canonical organization.

A promising line of network neuroscience, commonly referred to as Spectral Graph Theory, builds on the idea that structural connections might be the source upon which functional activity and behavior rely on [20, 21], although the degree of coupling between them is still being discussed [22]. Likewise, structural constraints can be used to model hemodynamics while potentially revealing effective connections [23]. Yet, the bonds between structural and functional connections remain, not surprisingly, an open problem [24, 25]. Unveiling mechanisms of structural plasticity evolution is therefore a key step towards understanding the impact of disruptions and recovery in both structural and functional connectomes. In this direction, and as a last contribution, we present a machine learning prediction study on how structural connections self-arrange after surgical resection of brain tumors.

Nonetheless, a certain number of challenges need to be addressed. Brain tumors, as well as their resections, critically affect elements of the network that may be far away from the damaged region itself [18]. Notwithstanding their success in representation learning and generative problems [26, 27], graph neural networks struggle with complex network topologies [28] due to difficulties in aggregating information from long-range connections. Dehmamy and colleagues showed how this can be bypassed by designing modular networks. However, this path inevitably leads to complex data-hungry architectures [29] that cannot be trained in sensitive clinical scenarios where data is scarce. Interestingly, fully connected layers, besides being easier to train, naturally combine knowledge regardless of neighbor proximity [28, 30, 31].

Thus, we build on the idea that healthy structural connections should be used to inform and guide predictions. We propose to use anatomical constraints in a Bayesian framework combined with fully connected layers to produce detailed graphs that share both visual similarities and network topological characteristics with the ground truth. In summary, our contribution is dual. We propose a prediction of structural rewiring post-surgery and we shed some lights on the impact of oedema in functional and structural signal.

## Results

### Tumor and Default Mode Network Functional Signals

We first addressed the question of whether functional signals were present inside the lesion. The segmented masks included all parts of the tumor, from the necrotic issue to the oedema (see Methods). For this study, resting-state fMRIs were available. We extracted functional signals from inside the lesioned parts of the brain and compared them with functional activity from the same region in control subjects (see Methods). Fast and naïve inspection of those signals did not yield visual differences between tumoral and healthy signals (Fig. 1A LEFT). As such, we qualitatively compared Blood-Oxygen-Level-Dependent (BOLD) signals from damaged tissues with signals from regions belonging to the Default Mode Network (DMN) (Fig. 1A RIGHT, see also Methods).

**Figure 1.**
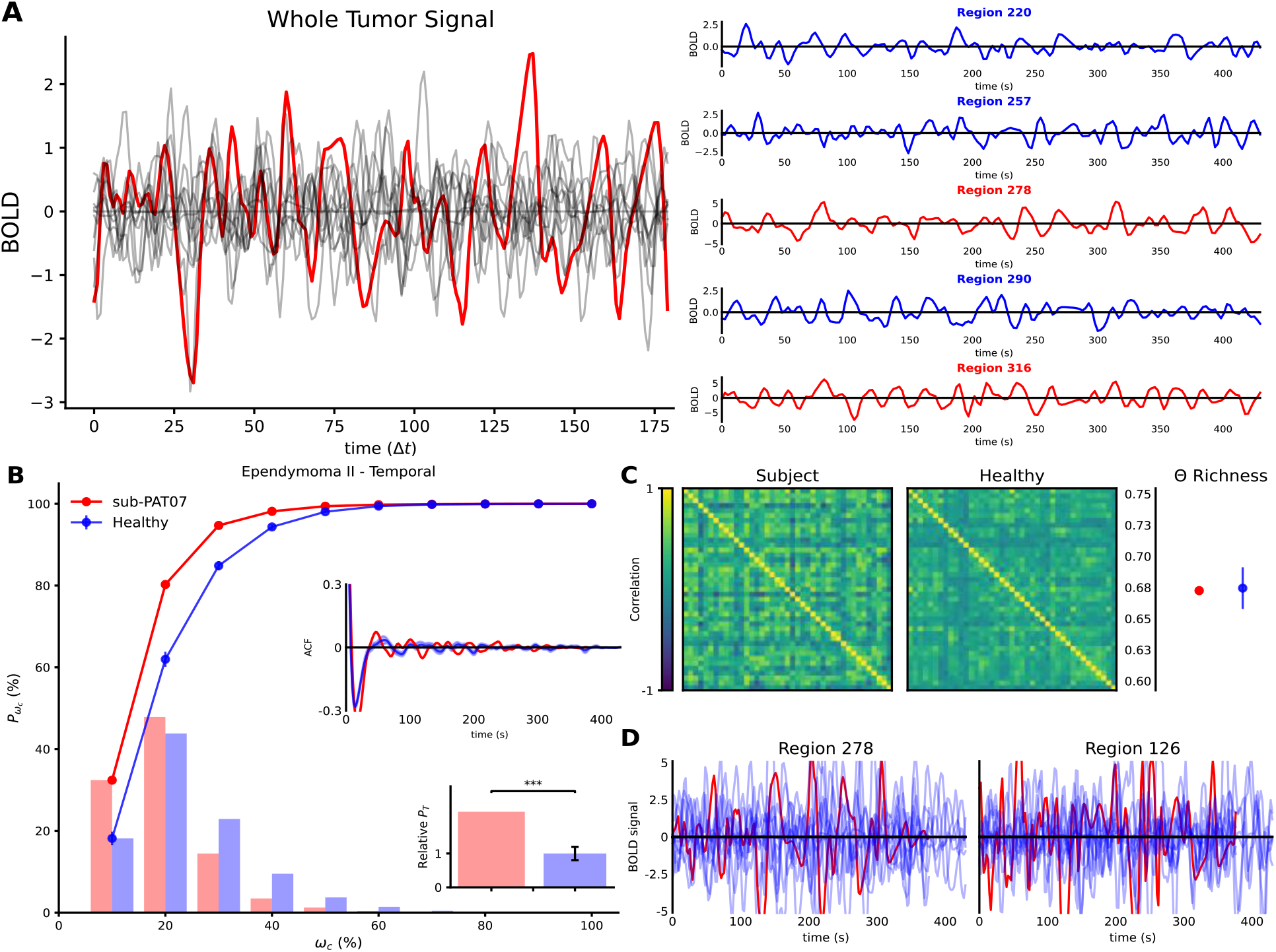
sub-PAT07 Oedema and DMN functional signals. **A)** LEFT Mean functional signal (BOLD) measured inside the oedema of a patient (red) as compared to signals from the same region in healthy subjects (gray). Similar signals were also found in all subjects inspected. Time axis is shown in time-steps rather than seconds. RIGHT Functional signals from 5 example regions belonging to the DMN of control subject 10 (randomly selected). Colors code for regions assigned to the same community through the Louvain Community detection algorithm. No qualitative difference is found between the raw time series inside the oedema of a patient and regions of the DMN of a healthy subject. **B)** Cumulative power (lines), power distribution (bars), autocorrelation function (ACF) and total power relative to healthy subjects (histogram) for the patient (red) and mean of controls (blue) functional signals of the regions in the DMN. Error bars [mean ± SEM] indiczate the results for the control population, while “*” code for statistical significance. **C)** LEFT Functional DMN from the same patient and mean of control subjects (see Methods). RIGHT Functional complexity as measured by the distribution of correlations for the patient’s network and the mean of control [mean ± SEM] (see Methods). **D)** Functional signal of two randomly selected regions from the same patient (red) as opposed to all the control subjects (light blue). No apparent difference is found between raw time series across regions, subjects and patients.

To further test whether functional activity inside tumors was related to global signals, we first studied how brain tumors themselves shaped those global signals. Direct comparison of averaged time series was not possible since arbitrary dephasings introduced artifacts (see Fig. S1A LEFT). Therefore, we analyzed functional series from regions belonging to the DMN in the frequency domain (see Methods). Alterations in the total power as well as the distribution of such power across frequencies were present in some subjects but not in others (Fig. 1B; see also Fig. S2-5A). The autocorrelation functions between time signals were more easily comparable, since they exhibited similar shapes for short time lags (Fig. 1B inset). We observed that left-skewed power distributions tended to higher autocorrelations than right-skewed power distributions (Fig. S4 inset) suggesting slower time series maintained temporal coherence for longer times. For longer lags, however, all signals lost this coherence. This inspired the definition of a score capable of distinguishing patients displaying faster oscillations from patients who showed the opposite. We expand on this idea in the next section.

A similar phenomenon was observed when pair-wise correlations between DMN regions were computed to reconstruct the network (Fig. 1C LEFT). Extensive research in network measures allowed us to estimate the complexity of networks based on the distribution of the correlations (see Methods). Briefly, we devised the Θ Richness score as the difference, in module, between the distribution of correlations building the network and a uniform distribution [32]. Differences in the Θ Richness between patients and healthy networks were also found to be inconsistent across subjects (Fig. 1C RIGHT; see also Fig. S2-5B). As a final step, we inspected signals from the same regions belonging to the DMN of both patients and control subjects, again finding no clear traces of tumoral damage in DMN functional signals (Fig. 1D).

In summary, functional signals from tumors and DMNs did not show significant changes in terms of complexity. However, they displayed alterations both in the power domain and in the distribution of functional connections.

### Temporal Dynamics and Default Mode Network Reorganization

To further characterize DMN signals in brain tumors bypassing phase and noise artifacts (perhaps unsuccessfully removed by the preprocessing pipeline), we designed a scalar score based on the cumulative power distribution of the time series. This Dynamics Alteration Score (DAS), inspired by previous work on periodic modelling of fMRI time series [33], was able to differentiate between slower or faster oscillations of BOLD signals (see Methods; see also Fig. S1). Time series dominated by slower oscillations with respect to healthy signals displayed positive DAS as more power was located to smaller frequencies in the spectrum (Fig. 1B). The opposite was found for a signal displaying faster oscillations (Fig. S4A).

Overall, the dynamics of DMN in patients was found to be positive, negative, or zero. However, visual differences between functional matrices (i.e., connectivity strengths) seemed to be anticorrelated with the magnitude of the DAS. We measured the similarity of the patients and control networks with a node similarity score (see Methods) and observed that it is significantly anti-correlated with the magnitude of the DAS (r=-0.39, p=0.027, one-tailed exact test; Fig. 2A LEFT).

**Figure 2.**
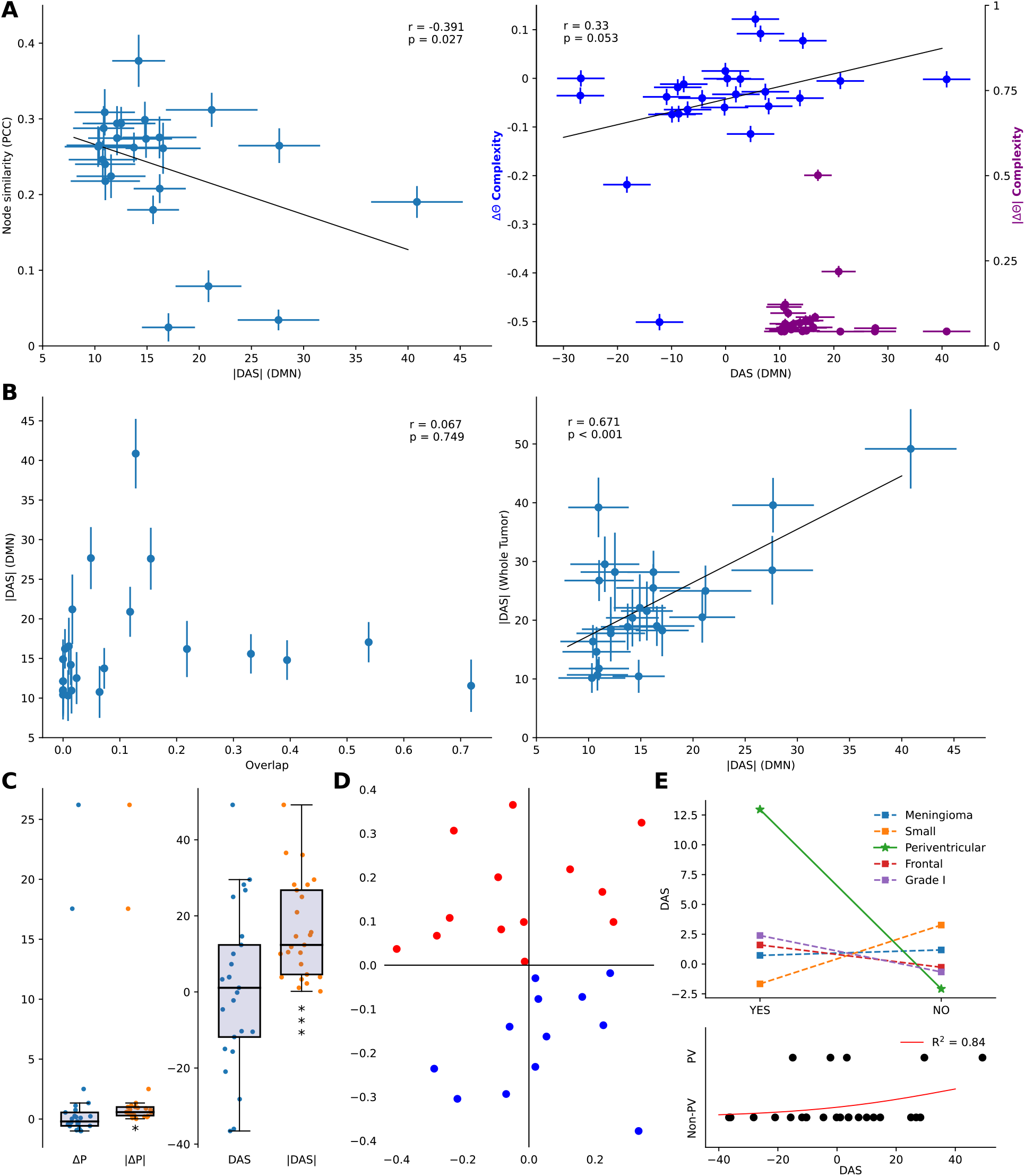
Linking DMN and Oedema BOLD Signals. Pearson Correlations and the corresponding p-values are shown. Error bars correspond to the mean ± SEM. **A)** LEFT Correlations between node similarity as measured by the Pearson correlation coefficient and the Dynamics Alteration Score (DAS). RIGHT Change in complexity of the patients’ DMN with respect to the healthy pool (Directionality ΔΘ and Magnitude |ΔΘ|). **B)** LEFT Scatter plot showing the null correlation between lesion overlap with DMN regions and the DAS. RIGHT Strong correlations existed between alterations inside the patients’ oedema and alteration in the DMN’s as measured by the DAS. **C)** Differences in total power relative to healthy subjects (LEFT) and dynamics (RIGHT). Only the absolute values where significantly different from zero (one-side t-test and U-test). **D)** Scatter plot of the two components found via Independent Component Analysis (FastICA) applied to the DAS between patients and control subjects. The same two clusters (red-blue) were consistently found with a K-means score of - 1.2. **E)** TOP Alteration in dynamics between different groups of tumors. BOTTOM Logistic regression between DAS and periventricular (PV) tumor patients. Qualitative inspection reveals a tendency for periventricular tumors to display slower BOLD dynamics, although they were found to be non-significant presumably due to the small number of samples (N=5, p=0.16, one-sided U-test).

Additionally, the magnitude of the change in Θ Richness between healthy and affected networks was found to be significantly greater than zero (p<0.001, one-tailed t-test). Opposite to this, the direction of this change was non-significantly different than zero (p=0.067, two-tailed t-test). Negative DAS implied the presence of higher frequencies of oscillations challenging the existence of coordinated oscillations. Interestingly, alteration on the dynamics showed a strong positive almost significant linear trend with the changes in functional complexity (r=0.33, p=0.052, one-tailed exact test; Fig. 2A RIGHT). Instead, when considering the magnitude of those changes, no statistical relationship arose (p=0.499, one-tailed exact test).

In conclusion, we found that changes in the signal oscillations of the DMN regions translated into significantly different patterns of functional connections. Crucially, the effects scaled with the magnitude and direction of the power distribution in the time series. In the following section, we explore how and where these alterations may arise in brain tumor patients.

### Local and Distributed Functional Signals Arising from Brain Tumors

Contrary to what we expected, the magnitude of alteration in the dynamics of DMN was not correlated with spatial overlap between the lesion and the corresponding DMN region (Fig. 2B LEFT, see also Methods). These results did not change when directly considering spatial distance (r=0.20, p=0.16, one-tailed exact test, see Methods). Next, to assess whether intra-tumor functional activity was both existent and relevant, we compared the DAS computed from the DMN with that obtained from intra-lesion signals (see Methods). The power of the signal from the tumor and the DMN was highly and significantly correlated (r=0.671 and p<0.001, two-tailed exact test; Fig. 2B RIGHT). This correlation was significant also when considering non-absolute values (r=0.677 and p<0.001, two-tailed exact test). This result ultimately linked what happened inside tumoral tissue with what was observed across spatially distributed brain regions belonging to a resting-state functional network.

We also compared the properties of the BOLD signal inside the tumor with the same regions in healthy subjects. The changes in total power were significant in magnitude (p=0.032, one-tailed t-test) but not in direction (p=0.36, two-tailed t-test; Fig. 2C LEFT). A similar result was found for the DAS, with the only difference of its effect being stronger (p<0.001, one-tailed t-test; Fig. 2C RIGHT), again similar to what we reported for the DMN regions.

Dimensionality reduction via Independent Component Analysis (FastICA) was applied to cluster differences between patients and control subjects. The same two clusters were consistently found with a K-means score of ∼-1.2, although no apparent groups were observable (Fig. 2D). As such, we hypothesized that the alteration in power distributions might be explained by tumor features. We divided subjects into two groups according to 5 tumor features: location (brain lobe), size (with respect to the sample median), periventricular (when located at the borders of the ventricles), histology (based on histological features of the resected tissue) and grade (based on the cell appearance and differentiation in the sampled tumor, which reflect tumor aggressiveness, i.e., the ability to grow, infiltrate and recur). The differences between the DAS group of whole tumors were rather high when considering whether the tumor intersected with the ventricles or not (Fig. 2E TOP), but this difference was not statistically significant (p=0.16, one-tailed U-test). However, given its larger magnitude and greater significance with respect to all other groups, we speculate on its meaning in the discussion. A logistic regression also displayed a reasonable determination coefficient and pattern (*R*^2^ = 0.84), but the small number of periventricular tumors did not allow for more quantitative assessment (Fig. 2E BOTTOM).

As a final note, we report that the tumor with the highest DAS was a grade II ependymoma located in the temporal lobe and intersecting with the left lateral ventricle. The tumor with the lowest (and negative) DAS corresponded with a grade I meningioma located in the frontal-skull base not intersecting the ventricles.

### Structural Brain Networks Containing Intra-tumoral fibers

As a first step towards reliable fiber tracking inside brain tumors, we designed a combined approach that used two different (previously validated) reconstruction algorithms (Fig. 3, see also Methods). However, validation of any fiber tracking pipeline is difficult due to the lack of any ground truth. As such, we compared the networks derived by the proposed approach against a widely used multi-shell pipeline.

**Figure 3.**
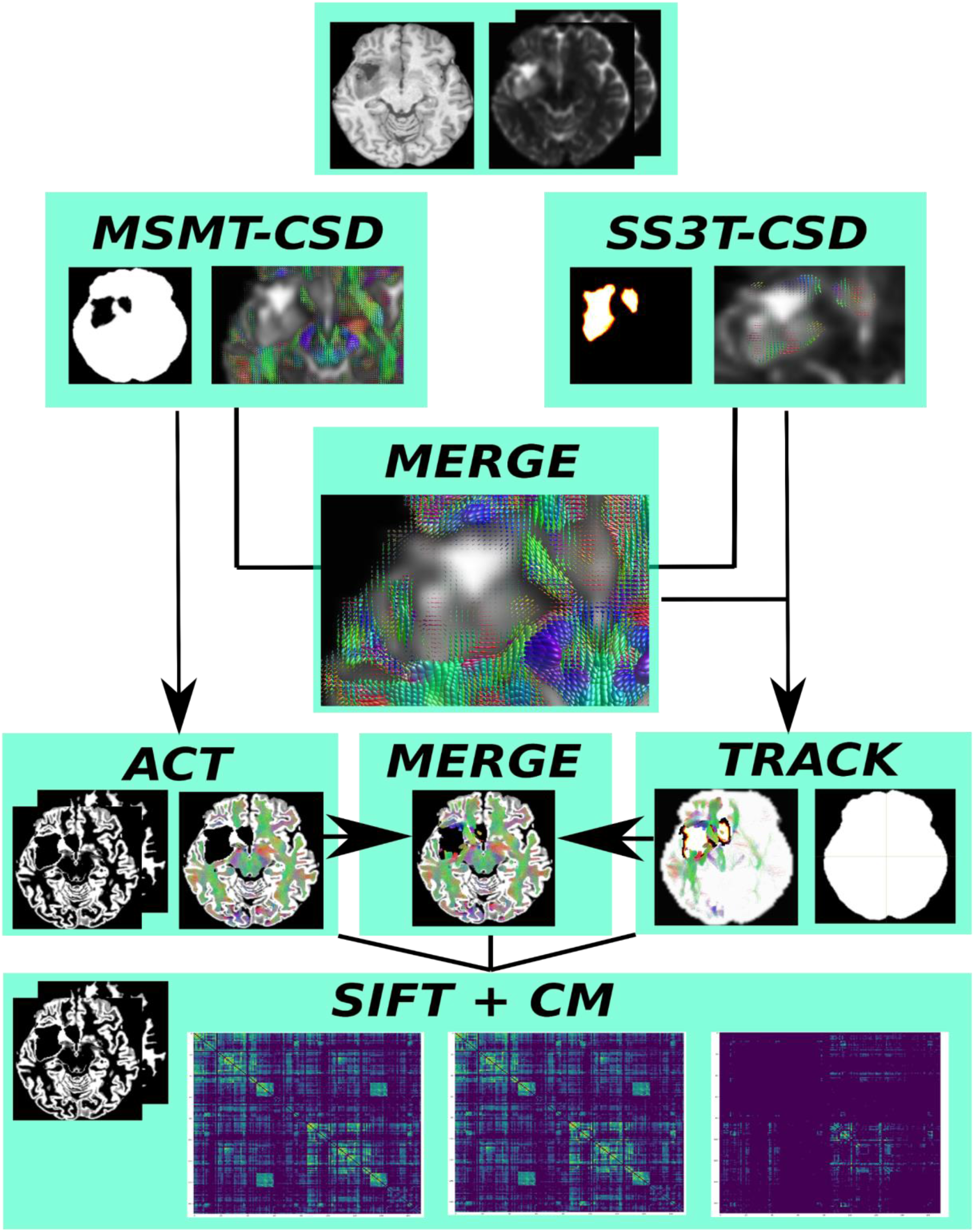
Hybrid reconstruction workflow. Besides the diffusion volumes, the pipeline requires as input a T1 weighted image from which a 5-tissue segmentation is derived. A binary mask is also extracted without including the. Multi-shell multi-tissue (MSMT-CSD) reconstruction methods are run together with anatomically constrained tractography (ACT) to obtain the “healthy” fibers and connections (bottom left connectivity matrix) without including lesioned tissue. Fiber orientation functions inside the oedemas are obtained by running the single-shell 3-tissue (SS3T) method only inside the lesioned regions (orange mask). Fiber orientation functions from both methods are then merged. Connections originating, terminating or traversing oedemic tissue are tracked with the iFOD2 probabilistic algorithm by only seeding inside the lesion. Maximum angle stopping criterion is used as an alternative for the anatomical constrain. Tractograms obtained with both branches are later combined using the *tckmerge* command from MRtrix3. To avoid double counting repeated streamlines, spherical-deconvolution informed filtering of tractograms (SIFT) is performed with the anatomical prior for the “healthy” and “merged” tractogram retaining 30% of the fibers. Eventually a symmetric connectivity matrix (CM) is obtained. In addition, a map of the connections traversing the oedema is also available after filtering streamlines without anatomical information (bottom right connectivity matrix).

We found that our workflow was able to maintain similar results as well as capture streamlines between brain regions that were lost when using the multi-shell algorithm alone (Fig. S5). This was expected since we specially set the pipeline to start streamlines inside the lesions. We also realized that in some tumors the Pearson Correlation Coefficient (PCC) between the patient network and the healthy cohort was increased with respect to the alternative pipeline while in some others not (Fig. S6B). Since brain tumors are heterogenous in many aspects, we tested if these differences were correlated to some features of the tumors, but we only found a very low, positive, non-significant correlation between differences in PCCs and tumor size (r=0.126 and p>0.55, exact test). Furthermore, a three-way ANOVA including tumor size, histology and grade revealed no significant relationships. However, the factor including the interaction between grade and size showed the smallest p-value (p=0.596).

### Structural Predictions After Tumor Resection

One of our main goals was to design a method capable of predicting how structural graphs will look after major surgical procedures while still elucidating the confounding effects of connectivity reorganization. For each reconstructed network, a total of 13695 normalized edges (see Supplementary Material) needed to be reconstructed, thus making the problem ill-posed (see Methods). Nonetheless, as argued in the introduction, we hypothesized that a fully connected network adequately guided with anatomical information could capture essential properties (both numerical and topological). We evaluated the model using Leave One Out Cross Validation therefore training and testing a total of 19 models or 19 *folds* (see Methods for training and testing details).

Previous work showed that linear models could surprisingly capture fundamental properties of structural graphs [34]. We evaluated a Fully Connected NETwork (FCNET) against a Huber Regressor and a null model. The choice of a Huber Regressor was motivated by its robustness to outliers and the presence of heterogeneous data points. Furthermore, testing against null models helps in avoiding circular analysis [35], therefore we also benchmarked FCNET against an untrained linear generator. Both the outputs of the Huber and ER generators were weighted by the same anatomical prior as the FCNET. For each model and fold, we tested the left-out network with 6 different metrics (see Methods). The results for each score are shown in Table 1 for the mean and in Table S1 for each subject in the dataset.

**Table 1.**
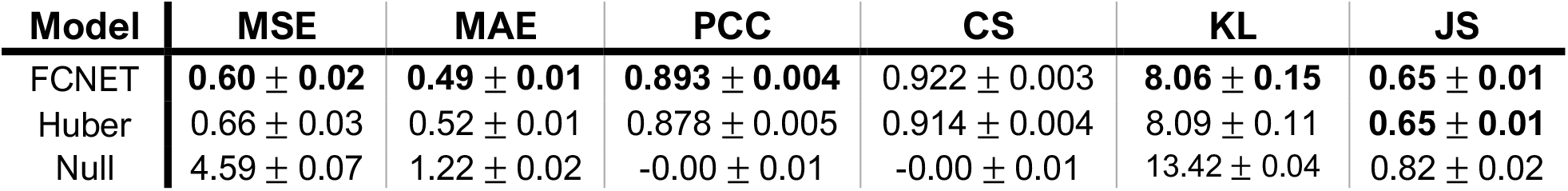
Model results (mean ± SEM). The Fully Connected NETwork (FCNET) was tested using a Leave One Out cross validation scheme in 6 metrics (MSE: Mean Squared Error; MAE: Mean Absolut Error; PCC: Pearson Correlation Coefficient; CS: Cosine Similarity; KL Kullback-Leiber and JS: Jensen-Shannon Divergences). In addition, we tested against two benchmark models: Huber and Null regressors. FCNET showed a significant improvement in all metrics evaluated. Both Huber and Null models were also tested in the same framework, that is, the predicted likelihoods weighted by the same anatomical prior (see Methods).

FCNET significantly outperforms the null model in all evaluation metrics (p<0.001, one-tailed t-test; p<0.001 one-tailed U-test). When tested against the Huber regressor, FCNET significantly outperformed it in all the metrics assessing numerical similarities (p<0.05, one-tailed t-test; p<0.05 one-tailed U-test). However, when tested for topological accuracy, FCNET did not improve with respect to the Huber regressor measured by the Kullback-Leiber (KL) and Jensen-Shanon (JS) divergences (p>0.31, one-tailed t-test; p=0.24, one-tailed U-test). Despite not being trained on preserving topological features, both FCNET and Huber captured structural properties since both models significantly decreased the KL (p<0.001, one way ANOVA; p<0.001 Kruskal-Wallis test) and JS (p<0.001, one way ANOVA; p<0.001 Kruskal-Wallis test) Divergences of the weight probability distributions between predicted and ground truth networks (see Methods).

None of the models was trained using a regularization method to prevent negative connections. Surprisingly, however, FCNET generated negative connections that accounted for less than 25% and they were all between 0 and -0.5. Since these values are in logscale they would account for a connection of less than 1 and probably getting filtrated by the anatomical threshold (see Methods). We show the generated post-surgery networks and residuals of three randomly selected subjects in Fig. 4.

**Figure 4.**
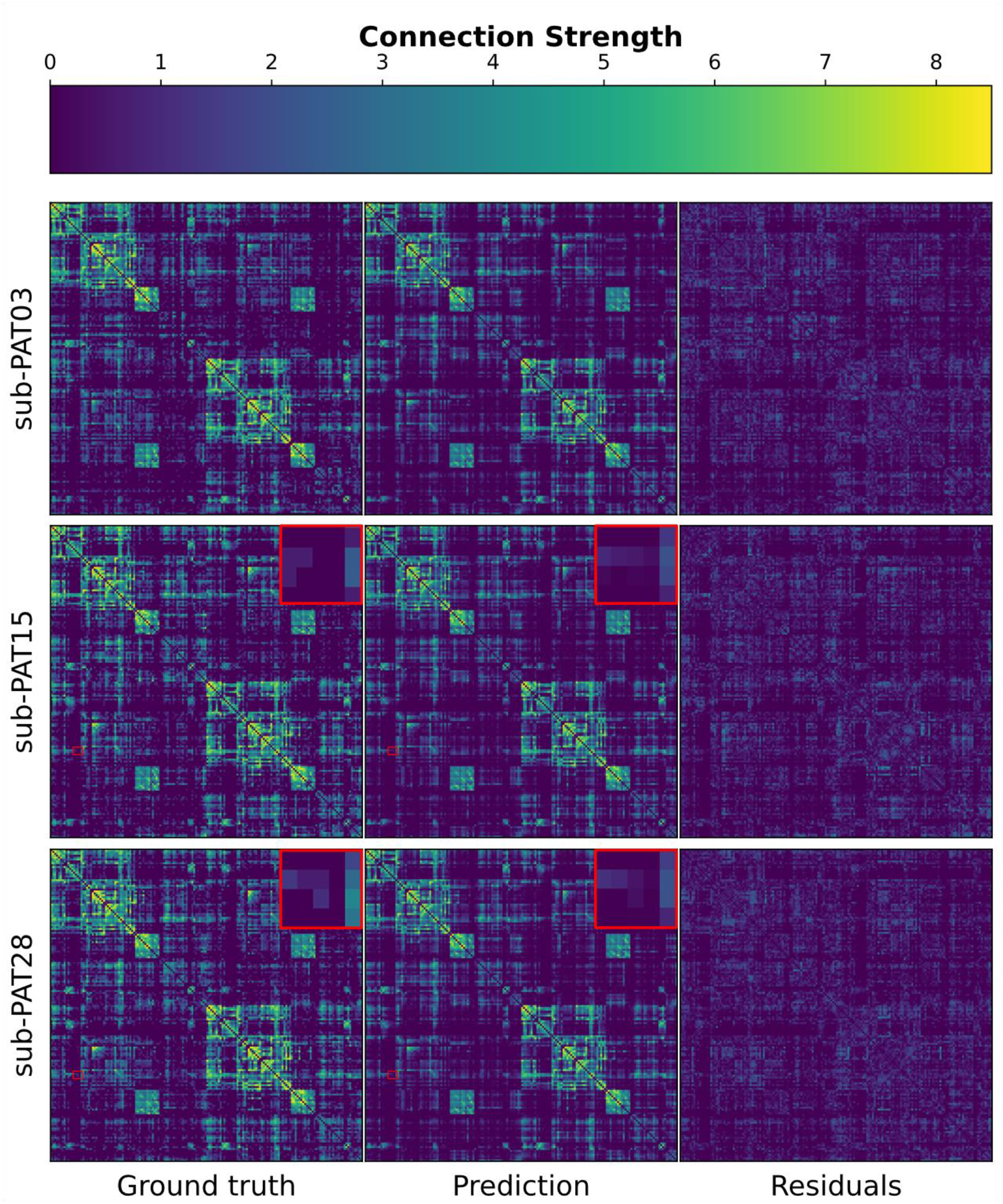
FCNET’s network generation. Three subjects were randomly selected to be displayed as a visual proof that FCNET captures essential properties of the post-surgery graphs. The residuals show the absolute difference between the predicted and ground truth networks. Negative connections were dropped for visualization purposes since they crucially affected the color scale but not the structure. FCNET can capture some specific inter-subject variabilities (augmented red squares) despite being trained on highly heterogeneous data. Connection strength is measured as *log*(1 + *w*) where *w* is the native connectivity derived from the tractograms.

### Subject-specific predictions

The connections within brain networks are highly heterogeneous in the context of a brain tumor. Herein, we imposed anatomical prior to act as a regularization method. However, a highly restrictive prior resulted in a complete loss of subject specificity despite FCNET achieving lower reconstruction errors. After some trial an error we found that an optimal (or nearly optimal) prior was able to discard enough connections while still capturing some inter-subject variability of the networks (Fig 4 red squares; see also Fig. S7 and Supplementary Materials). However, the model generalization does not allow for a perfect fit to the data, therefore a systematic error was present and observable in the residuals between the generated and ground truth networks (Fig 4 right column).

Next, we asked whether FCNET was simply overfitting a small subset of similar subjects. We calculated the z-score of each metric with respect to the 19 folds cross validated subjects (see Methods). For all metrics, we found that approximately 70% of all z-scores lied in the ±σ range and approximately 95% fell in the ±2σ. Even more, when repeating the training with different starting weights, all subjects but 2 showed different scores (Fig. 5 LEFT); they were not classified as outliers and therefore were considered in all the subsequent analysis (p>0.2 two-tailed Grubbs test). As a further checkpoint to ensure that the model was not overfitting to a specific subset of similar subjects (i.e., patients with similar tumors), we tested for the normality of the z-scores, thus finding that all of them had a significant linear correlation (r>0.98, p<0.001 one-tailed t-test) between the theoretical and observed quantiles (Fig. 5A RIGHT).

**Figure 5.**
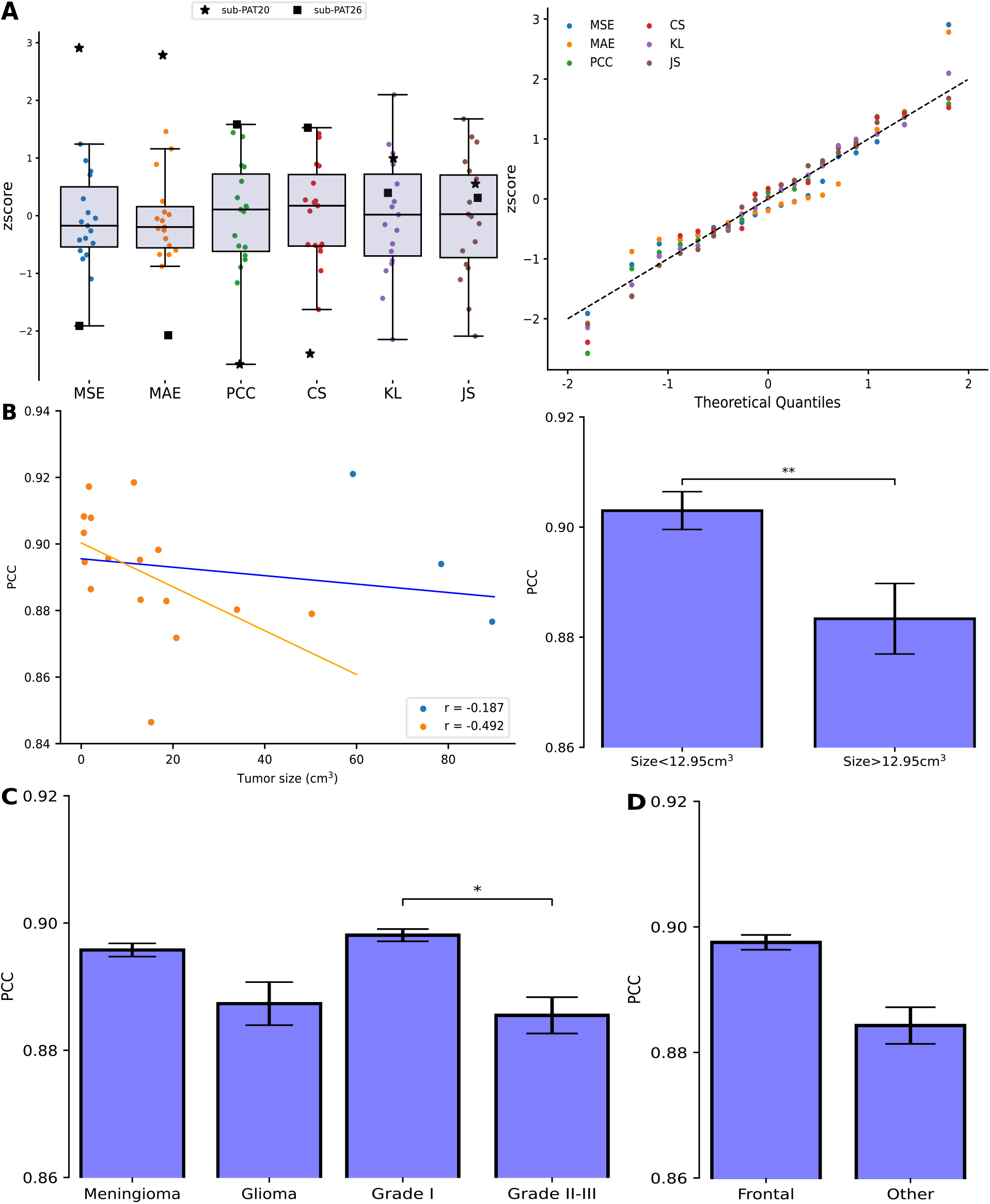
FCNET’s selective predictions. **A)** Normality assumptions on the whole pool of metrics used to test the model in each 1-fold (MSE: Mean Squared Error; MAE: Mean Absolut Error; PCC: Pearson Correlation Coefficient; CS: Cosine Similarity; KL Kullback-Leiber and JS: Jensen-Shannon Divergences). LEFT Boxplot of zscores. Shaded areas represent the Inter Quartile Range (Q3-Q1) and solid black line depicts the median. Marked subjects (PAT20 and PAT26) were not classified as outliers for all the metrics used (p∼0.15, Grubbs one-sided test). RIGHT Q-Q plot for each metric. Dashed black line shows the expected y=x relation. Significance for each regression was assed (r>0.95 and p<0.01). **B)** Impact of tumor size in the predicted graphs. LEFT Correlations between PCC and tumor size considering all but the three largest patients (orange; =0.011 one-sided t-test) and considering all subjects (orange+blue; p=0.251 one-sided t-test). RIGHT Differences in the PCC [mean ± SEM] between small and large tumors (Percentile 50). **C)** Differences in the PCC [mean ± SEM] between LEFT tumor type and RIGHT tumor grade (p=0.037, one sided U-test). **D)** Differences in the PCC [mean ± SEM] between tumor location.

Four metrics we used to evaluate the model’s output measured the numerical similarity between the ground truth and predictions. Not surprisingly, they exhibited high pair-wise correlations (|r|>0.98 and p<0.001, one-tailed permutation test), therefore post-hoc analysis on one score were generalizable to all. Unlike MSE and MAE, both the PCC and CS are normalized between -1 and 1 making it desirable regarding reproducibility. Without loss of generality, we based the following analysis on the PCC.

### FCNET’s Sensitivity to Tumor Size

Structural disconnections are strongly related to tumor size; thus, we hypothesized that large tumors would decrease the performance of the network. We found that prediction accuracy was sensitive to tumor size in the hypothesized manner. Considering all subjects, correlations between accuracy and size were rather strong (Fig.5B LEFT) but were not found to be statistically significant (r=-0.164, p=0.251, one-tailed t-test). To further investigate them, we redid the same analysis without the three largest tumors as they appeared to be abnormally large. Those tumors had volumes greater than 60 cm^3^ and their mean value was considered an outlier (p<0.01, one-tailed Grubbs Test). In this scenario, correlations drastically increased both in value and significance (r=-0.567, p=0.011, one-tailed t-test); yet, when disregarding the fourth largest tumor, relative changes in correlations were less marked (r=-0.630, p=0.010, one-tailed t-test).

To settle whether the effect of size was indeed present, we divided all patients, including the three largest ones, into two groups using the median of the whole dataset (Fig. 5B RIGHT). Subjects with small tumors (<12.95cm^3^) showed higher PCCs (0.902±0.004 [mean±SEM]) than subjects with large tumors (0.882±0.006 [mean±SEM]) (p<0.01, one-tailed t-test; p<0.01, one-tailed U-test).

In conclusion, the highly non-linear generative model was sensitively worse when considering large tumors. However, there seemed to exist confounding effects distorting the relationship. In the next section, we explored those in more detail.

### Potential Confounding Effects on Structural Connectivity Reorganization

The histology of the tumor greatly influences clinical considerations such as survival rates or possible treatment strategies. Meningiomas tend to exert pressure on the healthy brain tissues in opposite to gliomas which show more infiltrative behaviors. As such, white matter pathways might display different patterns of reorganization. However, we did not find significant differences (Fig. 5C LEFT; p=0.157, one-tailed t-test; p=0.098, one-tailed U-test) in the PCC between meningiomas (0.895±0.002 [mean±SEM]) and gliomas (0.886±0.003 [mean±SEM]). On the contrary, predictions were significantly better (p=0.037, one-tailed U-test) for low grade tumors (0.896±0.001 [mean±SEM]) than high grade (0.884±0.003 [mean±SEM]). We also noticed a non-significant effect according to parametric scores (p=0.11, one-tailed t-test), although given the rather limited sample size, its validity is limited.

Since not all regions of the brain are equally important in terms of structural connectivity, we also tested if the location of the tumor influenced had an impact on the predictions. The dataset included 12 patients with frontal and 7 patients with temporal or parietal tumors. We found nearly statistically significant differences in performance between frontal and non-frontal tumors (Fig. 5D). Despite not being significant (p=0.077, one-tailed t-test; p=0.059, one-tailed U-test), frontal tumors showed slightly higher PCC (0.896±0.002 [mean±SEM]) than temporal and parietal together (0.884±0.003 [mean±SEM]).

Lastly, and contrary to what we observed in functional dynamics, results of FCNET were not sensitive to whether the tumor intersected the ventricles (p=0.84, two-tailed U-test).

### Topological Predictions

We also tested the topology of the generated networks by computing the weight probability distribution. The loss function used to train all models did not have any topological term, but the generated networks shared global properties with the ground truth as measured by the KL and JS divergences in Table 1. The generated graphs showed a biological lognormal weight distribution with a small number of highly connected nodes (Fig. 6 LEFT; see also Supplementary Materials).

**Figure 6.**
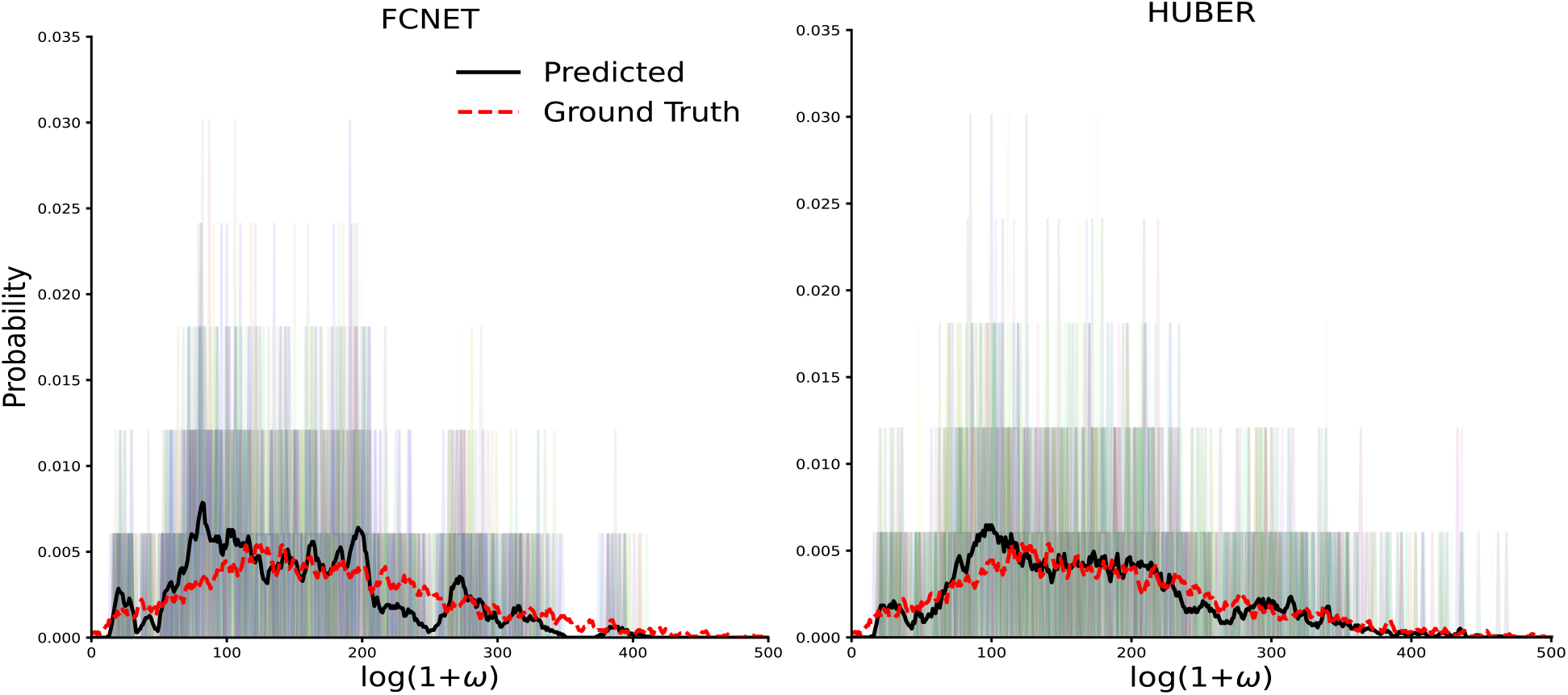
FCNET’s topological accuracy. Black thick lines show the mean weight probability distribution predicted by FCNET (LEFT) and the HUBER regressor (RIGHT). Dashed red line shows the mean weight probability distribution of the real post-surgery graphs. Shaded background bars show the predicted distributions for each subject in the dataset.

## Discussion

The focus of the present work is to quantitatively study functional and diffusion MRI signals in the presence of brain tumors. We show that when functional time series are compared in the frequency spectrum, the distribution of oscillatory frequencies is strongly related to how resting-state connections are re-wired. We also observe that the power spectrum within the brain tumor regions (including oedema) is strongly related to an oscillatory decoupling between the distributed and active brain regions. Moreover, we design a hybrid tractography pipeline capable of combining anatomical constraints and diffusion signal from the lesioned tissue. Finally, we use the reconstructed networks before and after brain surgery to train a machine learning model able to extract information about non-normative patterns of structural reorganization.

### Alterations in local oscillations are related to distributed functional desynchronization in brain tumor patients

We also set out to determine whether functional activity was present within the brain tumor. We find qualitatively similar signals between healthy and tumor tissues (Fig. 2A). However, the fact that there is a non-zero tumor functional signal is not indicative of its relevance. Therefore, to explore the functional sanity of the signals found, we first look at the existing differences in default mode networks (DMN) between patients and healthy controls.

Functional connections rely on temporal correlations between time signals which are entirely dependent on the phase as well as the active frequencies in the temporal signals [7]. Because of this mathematical construction, we hypothesized that the most descriptive feature of Blood-Oxygen-Level-Dependent (BOLD) time series would be the power distributions across frequencies. For this purpose, we designed a Dynamics Alteration Score (DAS) that can capture not only how different two power distributions are, but also how these differences are related with the relative dynamics of those signals (see Methods; see also Fig. 1B and S1-5). Intriguingly, the alteration in the signal dynamics (measured by the DAS), strongly correlated with network closeness (measured by node similarity; Fig. 2A LEFT).

The integrity of resting-state activity of the brain has been associated with cognitive function [6]. Based on this paradigm, numerous studies have explored how functional damage translates into cognitive impairment [4, 5, 1, 36, 3]. As a clear demonstration of the resilience of the brain to damage, all of them reported a continuous rather than binary decrease in cognitive performance. Brain networks undergo rearrangement to minimize the impact of lesions, and this rewiring follows specific constraints [24, 37]. From electroencephalography recordings, Wolthuis and colleagues found the level of topological similarity to be predictive of post-surgery cognitive skills [36], suggesting once again that understanding how the brain responds to lesions is crucial to predict the impact of surgical procedures on cognitive function.

To this purpose, we computed the complexity of the DMN and found significant alterations in magnitude with variable directionality across patients. Approximately half of the patients display increased network complexity while the rest exhibited the opposite pattern (Fig. 2A RIGHT). What these two non-contradictory findings suggest is that different tumors and/or patients react differently when considering DMN reorganization and inevitably lead to different network complexities. Curiously, a strong positive trend exists between the presence of slower oscillations (DAS> 0) and higher complexities (Fig. 2A RIGHT). We may speculate that faster time series resemble noisy temporal structures lacking temporal coherence, and therefore, resulting in a more uniform distribution of functional connections. In contrast, for slower functional dynamics, more complex networks would naturally arise. However, as stated by the proponents of the measure used here [32], high complexity does not imply an optimal segregation-integration balance promoting small-world topologies and their subsequent advantages in information propagation [38].

We also try to answer the origin of these desynchronizations. Current knowledge states that structural closeness is not sufficient to understand disruptions in patterns of functional disconnections [1, 2, 3]. In accordance with this, changes in BOLD dynamics are also not explainable by the spatial overlap nor distance between lesions and DMN (Fig. 2B LEFT). Instead, DMN desynchronization is better explained by frequency shifts inside brain tumors (Fig. 2B RIGHT). Current neuroscience methods struggle to address questions of causality, but for the sake of completeness and transparency of the results shown, we provide a speculative causal line to the findings reported. A plausible explanation would be to consider that the tumors themselves are altering the BOLD dynamics locally. Then, such an alteration would propagate through spatially distributed brain regions, perhaps supported by structural connections [23, 20, 22, 39], causing a global DMN desynchronization. To try and compensate, BOLD correlations between regions would rearrange themselves to spare the functionality. Network similarities and topologies across subjects would change, as found by previous studies [1, 2, 36, 3], without significantly altering their complexity. However, DASs could not disentangle whether the tumor altered the signal and that alteration spread to the DMN or the opposite.

Finally, periventricular tumors display a higher (positive) DAS than non-periventricular tumors. These differences do not reach statistical significance but are approximately five times larger in magnitude with respect to other groups. An increase in the volume of cerebral and lymphatic fluids inside the tumor may be the source of a larger desynchronization in the BOLD signal. Further research is required to determine whether this is associated with worse survival rates in higher grade glioblastomas [40, 41].

### Tracking fibers inside oedemic tissue

Brain networks, commonly referred to as connectomes, should be reconstructed with special care to avoid false positive connections. Moreover, in the presence of pathological tissue, state of the art fiber reconstruction methods suffer from cerebrospinal fluid contamination. Novel algorithms are specifically designed to minimize the effects of this contamination, but they neglect substantial information (i.e., multi-shell diffusion data) or are unable to apply anatomical constraints. In this regard, we designed a hybrid pipeline to use each method where it is best suited (Fig. 3). The resulting brain networks captured streamlines and connections which were previously non-existent when using each algorithm alone (Fig. S6A). Nonetheless, as ubiquitously seen in tractography studies, more streamlines do not necessarily mean biologically plausible fibers and/or connections.

Our proposed pipeline builds on top of a recently validated method. When used plainly, however, such an algorithm discarded multiple diffusion shells despite being usable outside the tumors [15]. On the other hand, Anatomically Constrained Tractography (ACT) has been shown to increase the correspondence between generated streamlines and real fiber bundles, but its usage is not suited for oedemic regions. Automatic segmentation algorithms often delineate tumors with gray matter causing premature stopping of tracking algorithms [19]. To bypass this effect, Aerts and colleagues [5] artificially copied the segmented brain tissue from the contra-lateral hemisphere to the damaged region. However, this strategy neglected the fact that a brain tumor could have critically altered the white and gray matter structures invalidating the proposed copied segmentation (Fig. S6A). In this direction, our pipeline used anatomical constraints outside tumors while entirely relying on the diffusion signal inside the tumor.

Alternative solutions to fiber tracking inside brain tumors include multicompartment diffusion tensor models but given their impossibility to resolve complex white matter structures as well as using diffusion signal from a single shell, we chose not to use them [13, 14].

### Anatomically informed generative model of brain structural connectivity after surgery

As a last contribution, with the reconstructed structural networks, we train a machine learning model to predict connectivity rearrangement after brain surgery. Brain tumors display high heterogeneity including size, location, histology, grade, and infiltration in gray matter areas, among others. Consequently, networks of brain tumor patients also show great variability (Fig. 4). Furthermore, interindividual variability in terms of brain plasticity and network rearrangement after surgery represents another relevant source of complexity when trying to understand and predict the organization of brain networks [42].

To reduce the impact of both factors, previous work [31, 43, 44, 45] guiding predictions in different context with networks from healthy subjects achieved good results, even when considering simple methods [34]. As such, we designed a flexible anatomical prior that was used to filter unplausible connections. We framed the problem in the Bayesian domain, which permits this prior to be backpropagated during the training phase (see Supplementary Material). Then, highly plausible connections are naturally given more importance when minimizing the loss function while, at the same time, successfully discarding improvable edges.

Neural architecture design in deep learning is itself an exciting and constantly evolving field. Nonetheless, it is well known how the choice of a specific architecture introduces a bias [46]. A natural choice for predicting brain connectivity is graph neural networks. However, as stated in the introduction they are not exempt of problems [28]. To overcome them, we design a one hidden layer non-linear regressor.

Fully connected layers have achieved great success in prediction studies of both clinical [47] and relational features [30], when adequately corrected for overfitting confounds. The model was trained with intercalated validation steps and its performance was evaluated using Leave One Out Cross Validation. The generative model achieved lower reconstruction errors than the chosen benchmarks (Table 1; see also Fig. 4). We also trained the same model using brain networks obtain without the hybrid tracking pipeline (see Methods). Interestingly, results were slightly worse, suggesting that our hybrid method is equivalent if not better than current state-of-the-art fiber reconstruction pipelines.

### FCNET surgical outcomes and aversive features for structural rewiring

Despite complying with normality assumptions, the reconstruction errors for two subjects were consistently low and high (Fig. 5A). An interesting paradigm in statistics emphasizes those data points that deviate from the trend. When looked from this perspective, the potential of deep learning models to uncover complex patterns translates into the increased ability to detect individual data points that deviate from, now more complex non-linear, statistical relationships.

The accuracy of the FCNET prediction was correlated with tumor size, achieving higher scores for small tumors (Fig 5B). The chances of affecting long distance structural connections naturally increase with the size of the tumor. Long distance connections are expected to carry high metabolic costs [37]. Moreover, long-range connections in structural networks have been shown to emerge during early stages of development in *C. elegans* [48], mice hippocampal circuits [49] and in primates [50], although debate is still active in the latter case. Therefore, directly reestablishing a long-range connection (i.e., end-to-end) to preserve rich-club and small-world topologies [5] might not be feasible. A possible way to circumvent these constraints would require several indirect connections leading to complex as well as non-normative patterns elusive to the model trained.

All but one patient suffered from grade I meningiomas, while all gliomas were of grades II and III. Interestingly, FCNET’s prediction accuracy is independent of tumor histology but not grade. According to previous studies, tumor grade and histology appear to have the greatest influence on survival rates in high-grade (III-IV) glioma patients [51]. Joint understanding of survival rates and connection rewiring is challenging, due to poorly stablished structure-function relationships. Given the results shown, however, we suggest that, for low grade tumors, grade instead of histology is a better indicator of normative patterns of connection reestablishment. In addition, the last years have been characterized by increasing evidence on the mutational profile impact of survival rate of gliomas [52]. This means that gliomas can have completely different survival rates according to their molecular profile [52, 53].

The location of the tumor did have an impact on the prediction score, although it was borderline significant. The predictions for frontal tumors showed slightly higher accuracies. Frontal areas are associated with higher metabolic activity [54] causing energy expenditure, shaping as well as incrementing the costs of network rewiring [55]. On the other hand, frontal regions are also the endpoints of fiber bundles, mainly involving both short- and long-range connections. Consequently, these two factors might compensate for each other, allowing for rather normalized patterns of network rewiring easily captured by the prediction model. However, it is difficult to assert the validity of this result as frontal tumors represented slightly more than half of the total. Thus, FCNET might very well be overfitting to these patients although we are inclined to discard this hypothesis given the precautions taken into consideration (Fig. 5A; see also Methods).

In contrast to the functional results discussed previously, the accuracy was independent of the periventricular features. This is understandable since the ventricles contain cerebrospinal fluid and no biologically relevant streamlines could be reconstructed within the ventricular system.

### Topologically meaningful predictions

Interestingly, despite not being trained on it, predictions from both the FCNET and the alternative model show essential topological properties found in brain networks. The predictive model generates weighted networks that, opposite to their simpler unweighted siblings, follow lognormal rather than scale-free distributions (Fig, 6; see also Supplementary Materials). This effect has been found in numerous studies on mice and macaques [56, 57], suggesting that rapid decay and high variance in connection strength arise from distance-dependent wiring costs [58].

### Limitations and future directions

Although we presented a validated tracking pipeline, more progress should be made in terms of filtering and identifying repeated connections derived by the merging strategy undertaken here (Fig. 3; see also Fig. S6). Instead of merging two tractograms, one could also merge the connectivity matrices resulting from them. This approach would also yield repeated connections; therefore, an ad hoc criterion should be carefully designed as to what connections should be kept or discarded.

However, the main limitation of this study is the small sample size and the lack of an external validation dataset. As such, all statistical results should be interpreted with extreme caution. Nonetheless, small sample sizes are found very often in studies dealing with patients with brain tumors. Even more, carefully designed metrics can also pinpoint existing phenomena despite not reaching statistical significance. Further work should also reveal the predictive potential of the score used in this work regarding cognitive impairment and perhaps local control and survival.

In terms of the machine learning model accuracy and reliability, due to a small sample size, we were limited as to which Deep Learning methods were usable. Recent progress in Geometric Deep Learning and Graph Representation Learning [26, 27] have yielded very promising results which are already showing great potential in medical imaging applications. Furthermore, it has been proven that topological guidance of graph neural networks drastically increases accuracy in highly heterogeneous data [29]. However, all these methods require huge datasets which may not be available for medically sensitive problems such as the one studied here. Even more, the structural topology and structure of networks suffering from brain tumors is not yet clearly understood, introducing a new layer of complexity as to what measures should be used to guide the training. Nonetheless, further work should find an optimal compromise to exploit these useful features in smaller datasets.

## Conclusions

Detailed analysis in the frequency domain revealed local and distributed abnormalities in resting-state time series and connections while stablishing a potential causal link between them. We also proposed a pipeline for fiber tracking that used more diffusion signal than previous attempts, as well as anatomical constraints. Lastly, we used these structural networks, that included intra-tumor streamlines and connections, to train a machine learning model to predict and study structural brain connectomes after surgery. The model achieved competent accuracies, disclosed tumor-dependent plasticity patterns, and preserved biological topologies. In summary, our results showed that brain tumors are both functionally and structurally dynamic, strengthening the need for more targeted MRI and neuroncology studies.

## Methods

### Acquisition and Usage of MRIs

A detailed explanation of the participants as well as the acquisition of the data is already available [4, 5]; nonetheless, for the sake of transparency we briefly present some crucial aspects. Subjects were asked to undergo MR scans both in pre- and post-surgery sessions. Out of the 36 subjects that agreed to take part in the pre-surgery session (11 healthy [58.6 ± 10.6 years], 14 meningioma [60.4 ± 12.3 years] and 11 glioma [47.5 ± 11.3 years]), 28 were scanned after a period spanning from 6 to 12 months in the post-surgery session (10 healthy [59.6 ± 10.3 years], 12 meningioma [57.9 ± 11.0 years] and 7 glioma [50.7± 11.7 years]). As a result, 19 pre- and post-surgery pairs of structural connectomes were usable as training and testing data. All brain tumors were classified as grade I, II, and III according to the World Health Organization.

Each MR session consisted of a T1-MPRAGE anatomical scan (160 slices, *TR* = 1750 ms, *TE* = 4.18 ms, field of view = 256 mm, flip angle = 9°, voxel size 1 × 1 × 1 mm^3^, acquisition time of 4:05 min) followed by a multi-shell HARDI acquisition (60 slices, *TR* = 8700 ms, *TE* = 110 ms, field of view = 240 mm, voxel size 2.5 × 2.5 × 2.5 mm^3^, acquisition time of 15:14 min, 101-102 directions b = 0, 700, 1200, 2800 s/mm^2^) together with two reversed phase-encoding b = 0 s/mm^2^ blips for the purpose of correcting susceptibility-induced distortions [59]. Resting-state functional echo-planar imaging data were obtained (42 slices, *TR =* 2100 ms, *TE =* 27 ms, field of view = 192 mm, flip angle = 90°, voxel size 3 × 3 × 3 mm^3^, acquisition time of 6:24 min). The TR was accidentally changed to 2400 ms after 4 control subjects, 5 meningioma patients and 2 glioma patients were scanned changing the times of acquisition to 7:19 min. For all the subsequent Fourier analysis, this time mismatch is solved by adding zero padding to the shorter time series.

Additionally, segmented lesions including the oedema, non-enhancing, enhancing and necrotic areas where available. Tumor masks were obtained with a combination of manual delineation, disconnectome [60] and the Unified Segmentation with Lesion toolbox^1^ [4].

### Pre-processing of MRIs

High resolution anatomical T1 weighted images were skull-stripped [61], corrected for bias field inhomogeneities [62], registered to MNI space [63] and segmented into 5 tissue-type images [64]. Diffusion weighted images suffer from many artifacts all of which were appropriately corrected. Images were also skull-stripped [61], corrected for susceptibility-induced distortions [59], denoised [65], freed from Gibbs ringing artifacts [66] and corrected for eddy-currents and motion artifacts [67]. The preprocessed images were then co-registered to its corresponding anatomical template (already in MNI space) [63], resampled to a 1.5 mm^3^ voxel size and eventually corrected for bias field inhomogeneities [62]. After motion correction as well as registration to the MNI template, the B-matrix was appropriately rotated [68].

Functional data was preprocessed with fMRIprep [69] and the eXtensible Connectivity Pipeline (XCP-D) [70] which are two BIDS compatible apps that perform all recommended processing steps to correct for distortion artifacts in functional data. Regression of global signal has been shown to improve denoising in BOLD series without excessive loss of community structure [71]. In total, 36 nuisance regressors were selected from the nuisance confound matrices of fMRIPrep output which included six motion parameters, global signal, the mean white matter, the mean CSF signal with their temporal derivatives, and the quadratic expansion of six motion parameters, tissues signals and their temporal derivatives [72]. Although smoothed time series were available, our analysis did not consider them. All specific steps were common to all subjects, both control and brain tumor patients. All images (T1s, T1 segmentations, diffusion, lesion masks and functional) were eventually co-registered to MNI space for comparison.

### Assessment of Default Mode Network and Oedemic BOLD Signals

BOLD signals belonging to the Default Mode Network were identified with the Gordon functional Parcellation [73]. More precisely, each one of the 41 regions classified as “Default” by the parcellation image was used as a binary mask to extract the time series from the functional image. For each subject (patient and control), the pair-wise Pearson correlation coefficient between time series was computed to obtain a functional connectivity matrix. The spatial overlap between DMNs and tumor masks was computed by summing all the voxels in the lesion mask belonging to one of these 41 regions. To normalize this score, we divided the resulting number by the number of voxels belonging to each one of the 41 regions labeled as “Default”. Note that, with this definition, an overlap of 1 would mean the presence of a tumor the size of the entire DMN.

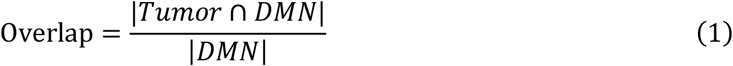

Moreover, the spatial distance between the center of mass tumor and the DMN was computed by averaging the Euclidean distances to center of mass of each one of the DMN nodes.

The DMN of the patients were compared to the mean of the healthy networks with two different metrics. Node similarity was assessed by computing the mean Pearson correlation between the same nodes in two different networks. Furthermore, to assess the complexity of a given network, we computed the absolute difference between the distribution of correlations building the network and a uniform distribution [32]. We refer to this score as Θ Richness:

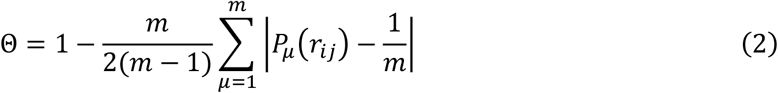

where *m* = 15 is the number of bins of the histogram estimating the distribution of correlations in the network *P*_*μ*_(*r*_*ij*_). Zamora-López and collueagues showed the robustness of the quantity in Eq. (2) with regards to the value of the parameter *m*. However, sensible choices range from 10 to 20 to ensure a sufficiently rich approximation of *P*_*μ*_(*r*_*ij*_). The changes in richness ΔΘ across patients were obtained by computing the difference relative to the richness of the mean DMN obtained from control subjects: ΔΘ = Θ_*Patient*_ – Θ_*Healthy*_.

A similar procedure was followed to study BOLD signals inside the lesioned tissue. For each patient, the binary mask containing the oedema was used to extract the time series from the patient, as well as from all control subjects. Consequently, BOLD signals in lesioned regions of the brain were comparable to 11 healthy signals from the same region. No network was computable in this case, making the use of Eq. (2) pointless.

### Fourier Analysis of BOLD Signals

To compare time series between subjects, we computed the Real Fast Fourier Transform of the BOLD series. This allowed us to compare the power spectrum of two or more signals regardless of, for example, the dephasing between them. Let *A*_*ω*_ be the amplitude of the component with frequency *ω*. Then, the total power of the signal can easily be obtained by summing the squared amplitudes of all the components:

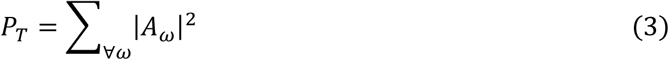

With the Fourier decomposition we could also characterize the power distribution of the signals as a function of the frequency. Analogous to Eq. (3), we summed the squared amplitudes corresponding to frequencies inside a bin of amplitude Δ*ω*.

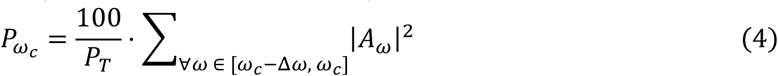

Since each signal had a different *P*_*T*_, to compare between subjects and/or regions, we divided the result by the total power *P*_*T*_ and multiplied by 100 to make it a percentage. Arbitrarily, we chose the parameter Δ*ω* for each subject so that each bin included 10% of the total power. The qualitative results did not depend on the exact choice of the bin width. Similarly, we computed the cumulative power distribution *CP*_*ω*_ by summing all the squared amplitude coefficients up to a certain threshold. For consistency, we measured the *CP*_*ω*_ as a percentage score and chose the thresholds to be multiples of exact percentages (i.e., *ω*′ ∝ 10%, 20%, …).

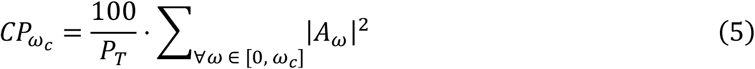

Both the power distribution *P*_*ω*_ and cumulative power distribution *CP*_*ω*_ can be used to compare dynamics between time series, but they have the inconvenience of not being scalar numbers. Furthermore, computing any distance-like metric (i.e., KL divergence) between these distributions across subjects would not yield any information of whether BOLD signals had slower dynamics (more power located in low frequencies) or the opposite (i.e., DMN in healthy and patient).

To overcome this, we designed a Dynamics Alteration Score (DAS) between time series based on the difference between two cumulative power distributions. It is worth noting that in the limit Δ*ω* → 0, the summations in Eqs. (2), (3) and (4) become integrals simplifying the following mathematical expressions. The DAS between two BOLD signals *i, j* was computed as the area between the two cumulative power distributions:

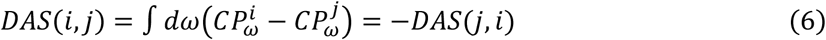

Finding a positive *DAS*(*i, j*) would mean that time series *i* had slower dynamics than time series *j* since more power is accumulated in lower frequencies with respect to the total. Throughout this manuscript, DASs were defined as the difference in power distribution between patients and the healthy cohort. For a simplified and, hopefully comprehensive, example, we kindly refer the reader to Fig. S1. To characterize a specific DMN, all these measures were computed for each region separately and then averaged [mean ± SEM]. As opposed to the Θ Richness, the DAS was computable both for DMNs and tumors since it only required two temporal series rather than a complete distribution.

For the score defined in Eq. (6) to make sense, the Real Fast Fourier Transform of the time series needed to be computed using the same frequency intervals, which, in short, implied that the time duration of the signals needed to be equal. For functional images with different *TR*s, this was solved by adding zero-padding to the shortest signal to match the same time duration. Nonetheless, this correction was added later, and its effect was found to be qualitatively and quantitatively non-existent.

### Reconstruction of Oedemic Structural Brain Networks

To ensure a detailed subject-specific network, we used a state-of-the-art pipeline to obtain brain graphs while at the same time not neglecting tracts inside lesioned regions of the brain (i.e., brain tumor oedemas). We combined two reconstruction methods, yielding three different tractograms and connectivity matrix for each one of them. A schematic workflow of the pipeline is in Fig. 1.

The first branch of the method consisted of a well validated set of steps to reconstruct the network without considering lesioned regions of the brain. For each b-value shell and tissue type (white matter, gray matter and cerebrospinal fluid) a response function was estimated [74]. The fiber orientation distribution functions were built and intensity normalized using a multi-shell multi-tissue constrained spherical deconvolution approach [75]. Anatomically constrained probabilistic tractography (with 10M seeds,4M streamlines, cutoff value of 0.08, min/max length of 3/280 and a step size of 0.75mm) was performed using dynamic seeding [64] and the iFOD2 algorithm [76]. To further improve the correspondence between the tractograms and preprocessed DWIs, we used spherical-deconvolution informed filtering [77] retaining a total of 1.2M streamlines (30%). To make sure that the lesioned tissue was not being considered, we used a binary brain mask that did not include the segmented lesion. This step was added for consistency with the logic of not tracking within the oedema. Nonetheless, the steps were repeated without this mask and the results were found to be almost identical. This was expected as multi-shell methods highly disregard cerebrospinal fluid contamination inside the lesion.

Next, to consider fiber bundles that originate and traverse oedemic tissue, a recent method for reconstruction was used only in the segmented lesion [17]. The coined Single-Shell-3-Tissue Constrained Spherical Deconvolution (SS3T-CSD) algorithm uses only one diffusion shell and the unweighted b=0 volumes. We used the b=2800 s/mm^2^ shell as it allowed for a better response function estimation for all 3 tissues [74]. Furthermore, to include long-range connections that might originate in the damaged tissue, we merged the reconstructed fiber orientation functions together with the previously obtained with the multi-shell approach (Fig. 1CENTER). It is important to note that both images were in NIFTI format and were already co-registered, therefore making this step straightforward. As already mentioned in the main text, anatomical constraints are no longer suited since white and gray matter are heavily compromised inside the lesion. Even more, the anatomical constrain caused fibers to stop around the oedema since those surrounding regions were (nearly-)always segmented as gray matter. Dynamic seeding was applied only within the masked lesion and a maximum angle of 0.08 was set as stopping criterion for the iFOD2 probabilistic tracking algorithm. Each brain tumor had very different features including size and cerebrospinal fluid contamination making the choice of the number of seeds and streamlines difficult (if not useless). For consistency, these numbers were kept at 10M for the seeds and 4M for the streamlines, although they were never reached because the volume of the lesion did not allow it. Spherical-deconvolution informed filtering was applied without anatomical information. As a result, a tractogram only containing fibers originating, terminating, or traversing in the lesioned brain regions. Once again, the percentage of filtration was difficult to determine, so the default values of MRtrix 3 were kept [78]. Other choices were tested for these two last steps, but the results were not affected by the choice.

The final and main branch of the method consisted in merging the two tractographies using the *tckmerge* command available in MRtrix3 [78]. Crucially, a streamline originating within the oedema might be the same as that originating in the surrounding regions. Consequently, merging both tractogram files would introduce false positive connections. To correct for this, spherical-deconvolution informed filtering was performed with anatomical information encoded in the 5-tissue-type segmented image retaining once again approximately 1.2M (30%) streamlines. It is worth noting that the number of reconstructed streamlines from lesion masks was about 2 orders of magnitude less than whole brain tractograms.

All resulting tractograms were then mapped to the third version of the Automated Anatomical Labeled atlas [79] to obtain a symmetric connectivity matrix between 166 brain regions for each subject^2^. Since a total of 3 tractograms were computed, 3 different connectivity matrices were available for comparison and further studies if needed (Fig. 1BOTTOM): the full connectome, the “healthy” connectome without oedemic connections and the connections originating/terminating or traversing tumor regions. The networks from control subjects and post-operative patients were reconstructed using the usual multi-shell multi-tissue branch without the binary lesion-free mask. Alternatively, networks from pre-operative patients were reconstructed with the described hybrid pipeline.

Track-density maps were computed from the tractograms and T1 weighted images with the *tckmap* command available in the MRtrix3 software package.

### Anatomical Priors and Bayesian Filtration of Predictions

As suggested by previous works, guiding learning with healthy cohorts should be useful to inform predictions [43, 44, 31]. Brain graphs are notoriously heterogeneous when considering age related differences. To take this into account, we selected subjects with significant age overlap between healthy subjects and patients in both tumor types. However, we did not consider sex segregation, since structural differences are rather unclear [80]; even more, the sample size for each subgroup would be severely reduced. We built a prior probability distribution of healthy links to guide the predictions using a *thresholded* average of the set of connections present in this healthy cohort (see Supplementary Material). This thresholded average allowed us to control both for the inclusion (or exclusion) of spurious connections and minimizing the false positive rate of connections [81].

### Training and Testing the Machine Learning Model

The high number of reconstructed fibers yielded high values for the connectivity between ROIs (∼10^3^). To prevent numerical overflow as well as to enhance differences in lower connections, all weights *w* were normalized by computing *log*(1 + *ω*) before feeding them into the artificial deep neural network.

The model consisted of a 1 hidden layer deep neural network which was trained minimizing the Mean Squared Error (MSE) between the output and the ground truth determined from the MRIs (see Supplementary Material). The weights were optimized using stochastic gradient descent with a learning rate of 0.01 and 100 epochs to avoid overfitting. Evaluation metrics included the Mean Absolute Error (MAE), Pearson Correlation Coefficient (PCC) and the Cosine Similarity (CS) between the flattened predicted and ground truth graphs. The topology of the generated networks was evaluated by computing the Kullback-Leiber and Jensen-Shannon divergences between the weight probability distributions of the generated and real graphs.

Leave One Out cross validation was done using 18 connectomes to train each one of the 19 models. For each model, the training data was randomly split into train (80%) and validation (20%) sets to prevent overfitting. Validation steps were run every 20 training epochs. For each fold, the testing of each model was done in the *left-out* connectome (Table S1).

### Statistical and Topological network analysis

Statistical tests and p-value computations were done with Scipy’s stats module. Unless stated otherwise, we use one-tailed approaches when addressing significance of strictly positive magnitudes (i.e., absolute values) or increases and decreases of performance. When testing for the direction of a certain result, we run two-tailed tests.

The Leave One Out cross validation approach yielded a pool of 19 subjects that were independently tested. For each metric, we computed the z-score by subtracting the mean and dividing by the standard deviation of the sample. Despite verifying that all of them were normally distributed, we run parametric and non-parametric statistical tests due to the small sample size. Topological metrics were computed using the Networkx python library [82].

Since the brain graphs were weighted, we computed a weight probability distribution instead of the more common degree distribution (see Supplementary Material). To compare the weight probability distributions of two graphs, we computed the Kullback-Leiber as well as the Jensen-Shannon divergences. The Jensen-Shannon divergence has the advantage of being both symmetric and normalized between 0 and 1 therefore interpretable as a distance between two distributions (i.e., predicted vs ground truth).

## Acknowledgements

J.F.-R. and A.Cr. thank Sofie Valk and Daniel S. Marguiles for extremely helpful discussions about functional data and connectivity. This research was supported by European Union’s Horizon 2020 research and innovation program under grant agreement Sano No 857533 and by Sano project carried out within the International Research Agendas program of the Foundation for Polish Science, co-financed by the European Union under the European Regional Development Fund. The authors declare no other competing interests.

## Author Contributions

J. F.-R. and A.Cr. conceptualized the study; J. F.-R. analyzed the data; J.F.-R., F.S., A.Ca. and A.Cr. wrote the paper.

## Data and Code Availability

The original data is publicly available at OpenNeuro [60]. For the processing of MRIs several software packages were combined and integrated in a flexible python 3.9.7 pipeline [69, 70, 78, 83, 84, 85]. For the machine learning model PyTorch and CUDA libraries were used. Full disclosure via an [Open Science Framework](https://osf.io/48fnm/) repository^3^.

## Supplementary Material

### Generation of post-surgery graphs with artificial neural networks

A connectome for the *n*-th subject at a particular time point *t* which, in our case, was mapped to control, pre- and post-surgery stages *t*_*c*_, *t*_*pre*_ and *t*_*post*_. Only the lower triangular part of the connectomes was flattened into a vector ***x***_*n*_(*t*) of *E* components, given that structural connections are always symmetric and no self-loops were considered. Each one of these components *x*_*nk*_(*t*) with *k* = 1, …, *E* represents a link of strength *ϵ* between two brain regions. We simplify the notation and refer to pre-surgery connectomes as ***x***_*n*_, to post-surgery connectomes as ***y***_*n*_ and to healthy connectomes as ***z***_*n*_.

Based on previous work [1], we defined a distribution of binary links based on connectomes from *N*_*C*_ healthy subjects. For the *k*-th edge, *λ*_*k*_ = 0,1 is a Bernoulli distributed binary variable with probability

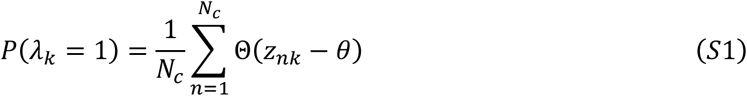

where Θ(·) is the Heaviside step function and *θ* = 0.2 is a threshold that filters links with insufficient strength and therefore minimizing the false positive rate. For the choice of the threshold, we considered the Shannon entropy and the modularity [1, 2] of network built according to Eq. (S1). We observed that small thresholds maintained a high entropy allowing for the inclusion of more connections without excessively altering the structure of the prior (Fig. S7). Lower thresholds allow more variability in the generated prior since more spurious connections can be considered.

For each patient and edge, the post-surgery probability of having a meaningful link *λ*_*k*_ = 1 with strength *ϵ* is conditioned on the pre-operative graph. We exploit this fact with the well-known Bayes’ theorem

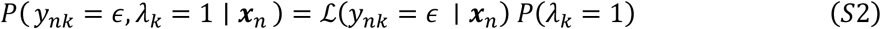

where ℒ( *y*_*nk*_ = *ϵ* | ***x***_*n*_) is the likelihood of having a post-surgery *ϵ*-strengthened connection conditioned on the pre-surgery connectome. For Eq. (S2) to hold, *P*(λ_*k*_ = 1) must be independent on the pre-surgery graph ***x***_*n*_. This assumption is reasonable since the anatomical prior was built using only healthy controls, therefore, not considering lesioned connections. We sampled each *k*-th connection using the Maximum A Posteriori criterion to generate the post-surgery graphs. To train the network, the sampled connections were then compared to the ground truth using the Mean Squared Error (MSE).

Although many possibilities emerge for estimating the likelihood in Eq. (S2), we used a fully connected network with one hidden layer (linear plus a sigmoid activation function). Several options were considered here, but we found that adding more layers or even considering 1D or 2D convolutions did not add significant improvements to the model. We report the mean results for a two hidden layer fully connected layer in Table S2.

### Weight probability distribution of brain networks

For all graphs, we computed the degree of each node by summing over all the connections

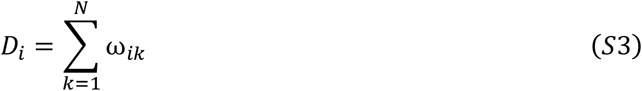

where *N* is the number of nodes, 166 in our graphs. To compute the probability distribution of a node having degree *d*, we counted all the nodes whose degree fell in a bin of width Δω as

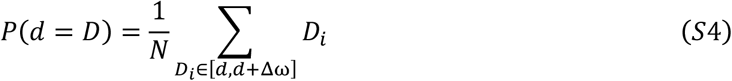

where *N* is again the number of nodes that acts as a normalization constant.

Networks wired at random have a binomial degree distribution, however it was shown that biological networks deviate from this tendency towards a scale-free or, in the specific case of brain networks, a broad-scale distributions [3, 4]. In both cases, the number of highly connected nodes (i.e., nodes with large degree) decays following a power-law in the case of scale-free and an exponentially truncated power-law for brad-scale distributions. However, for weighted networks, it has been shown that a lognormal distribution is better suited to describe brain connections [5, 6].The probability density of a lognormally distributed variable follows

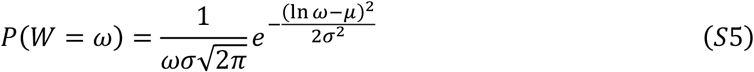

where *μ, σ* are the geometric mean and standard deviation. Note, that the main manuscript deals with the variable 1 + *ω*, therefore Eq. (S5) should be corrected for this. Given that the Jacobian of the transformation is 1, the proof is straightforward.

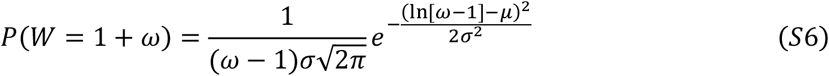

Therefore, the variable 1 + *ω* also follows a lognormal distribution as seen in Fig. 6 of the main text.

### Subject specific results and model comparison

**Table S1.** Leave One Out cross validation results and a hybrid fiber reconstruction pipeline. Results for the tested subject in each trained model. (MSE: Mean Squared Error; MAE: Mean Absolut Error; PCC: Pearson Correlation Coefficient; CS: Cosine Similarity; KL Kullback-Leiber and JS: Jensen-Shannon Divergences). For each fold the subject tested was not included in the training set (see Methods). The best subject in each according to each metric is highlighted in bold. Correlations between all pairs of metrics were high (r>0.95 and p<0.01). Only in PCC and CS a higher number means a better fit.

| Subject | MSE | MAE | PCC | CS | KL | JS |
| --- | --- | --- | --- | --- | --- | --- |
| sub-PAT25 | 0.657 | 0.495 | 0.883 | 0.916 | 8.07 | 0.648 |
| sub-PAT05 | 0.601 | 0.486 | 0.883 | 0.917 | 8.42 | 0.660 |
| sub-PAT11 | 0.662 | 0.500 | 0.872 | 0.901 | **6.67** | **0.602** |
| sub-PAT28 | 0.552 | 0.483 | 0.918 | 0.943 | 8.16 | 0.664 |
| sub-PAT08 | 0.572 | 0.470 | 0.898 | 0.923 | 7.89 | 0.634 |
| sub-PAT15 | 0.596 | 0.489 | 0.908 | 0.936 | 8.76 | 0.668 |
| sub-PAT17 | 0.591 | 0.482 | 0.903 | 0.931 | 7.56 | 0.647 |
| sub-PAT07 | 0.707 | 0.534 | 0.874 | 0.912 | 9.42 | 0.684 |
| sub-PAT24 | 0.575 | 0.481 | 0.896 | 0.927 | 7.66 | 0.628 |
| sub-PAT06 | 0.558 | 0.487 | 0.895 | 0.927 | 7.75 | 0.638 |
| sub-PAT02 | 0.568 | 0.472 | 0.895 | 0.925 | 8.22 | 0.652 |
| sub-PAT10 | 0.548 | 0.464 | 0.908 | 0.935 | 7.52 | 0.629 |
| sub-PAT01 | 0.522 | 0.470 | 0.917 | 0.942 | 7.96 | 0.644 |
| sub-PAT26 | **0.451** | **0.430** | **0.921** | **0.944** | 8.32 | 0.654 |
| sub-PAT03 | 0.608 | 0.512 | 0.894 | 0.926 | 8.86 | 0.657 |
| sub-PAT20 | 0.822 | 0.564 | 0.846 | 0.890 | 8.70 | 0.659 |
| sub-PAT23 | 0.626 | 0.477 | 0.877 | 0.910 | 7.44 | 0.623 |
| sub-PAT13 | 0.584 | 0.474 | 0.886 | 0.916 | 7.13 | 0.612 |
| sub-PAT16 | 0.697 | 0.528 | 0.879 | 0.915 | 8.64 | 0.677 |

**Table S2.** Comparisson of alternative models (mean ± SEM). Comparisson of alternative models using a Leave One Out cross validation scheme in 6 metrics (MSE: Mean Squared Error; MAE: Mean Absolut Error; PCC: Pearson Correlation Coefficient; CS: Cosine Similarity; KL Kullback-Leiber and JS: Jensen-Shannon Divergences). Last row corresponds to the same FCNET model but using as inputs brain networks obtained with the alternative multi-shell multi-tissue fiber reconstruction pipeline (see Main Text; see also Methods).

| Model | MSE | MAE | PCC | CS | KL | JS |
| --- | --- | --- | --- | --- | --- | --- |
| FCNET | **$0.60 \pm 0.02$** | **$0.49 \pm 0.01$** | **$0.893 \pm 0.004$** | $0.922 \pm 0.003$ | **$8.06 \pm 0.15$** | **$0.65 \pm 0.01$** |
| 2-layer | $0.68 \pm 0.02$ | $0.52 \pm 0.01$ | $0.892 \pm 0.004$ | **$0.923 \pm 0.004$** | $8.19 \pm 0.14$ | **$0.65 \pm 0.01$** |
| FCNET<br>(*msmt*) | $0.62 \pm 0.02$ | $0.49 \pm 0.01$ | $0.892 \pm 0.004$ | $0.922 \pm 0.003$ | $8.23 \pm 0.12$ | $0.66 \pm 0.01$ |

**Table S3.** Leave One Out cross validation results and a multi-shell multi-tissue fiber reconstruction pipeline. Results for the tested subject in each trained model when using only a multi-shell multi-tissue fiber reconstruction method. (MSE: Mean Squared Error; MAE: Mean Absolut Error; PCC: Pearson Correlation Coefficient; CS: Cosine Similarity; KL Kullback-Leiber and JS: Jensen-Shannon Divergences). For each fold the subject tested was not included in the training set (see Methods). The best subject in each according to each metric is highlighted in bold. Correlations between all pairs of metrics were high (r>0.95 and p<0.01). Only in PCC and CS a higher number means a better fit.

| Subject | MSE | MAE | PCC | CS | KL | JS |
| --- | --- | --- | --- | --- | --- | --- |
| sub-PAT25 | 0.694 | 0.509 | 0.875 | 0.911 | 8.03 | 0.596 |
| sub-PAT05 | 0.623 | 0.496 | 0.878 | 0.913 | 8.23 | 0.633 |
| sub-PAT11 | 0.641 | 0.482 | 0.876 | 0.904 | 7.53 | 0.567 |
| sub-PAT28 | 0.548 | 0.483 | 0.920 | 0.944 | 9.18 | 0.630 |
| sub-PAT08 | 0.613 | 0.486 | 0.889 | 0.921 | **7.07** | 0.601 |
| sub-PAT15 | 0.619 | 0.499 | 0.905 | 0.934 | 8.70 | 0.611 |
| sub-PAT17 | 0.598 | 0.487 | 0.900 | 0.929 | 8.76 | 0.602 |
| sub-PAT07 | 0.707 | 0.534 | 0.874 | 0.912 | 7.96 | 0.652 |
| sub-PAT24 | 0.549 | 0.472 | 0.900 | 0.931 | 8.00 | 0.639 |
| sub-PAT06 | 0.600 | 0.500 | 0.886 | 0.920 | 8.11 | 0.592 |
| sub-PAT02 | 0.570 | 0.471 | 0.895 | 0.925 | 7.82 | 0.649 |
| sub-PAT10 | 0.564 | 0.473 | 0.903 | 0.932 | 9.28 | 0.627 |
| sub-PAT01 | 0.506 | 0.462 | 0.918 | 0.943 | 8.32 | 0.641 |
| sub-PAT26 | **0.459** | **0.430** | **0.921** | **0.945** | 8.21 | 0.601 |
| sub-PAT03 | 0.605 | 0.511 | 0.893 | 0.925 | 8.14 | 0.657 |
| sub-PAT20 | 0.780 | 0.549 | 0.855 | 0.896 | 8.44 | 0.641 |
| sub-PAT23 | 0.611 | 0.469 | 0.880 | 0.913 | 7.55 | 0.610 |
| sub-PAT13 | 0.564 | 0.466 | 0.891 | 0.919 | 8.74 | 0.605 |
| sub-PAT16 | 0.698 | 0.529 | 0.877 | 0.914 | 8.29 | **0.560** |

---

**Figure S1.**
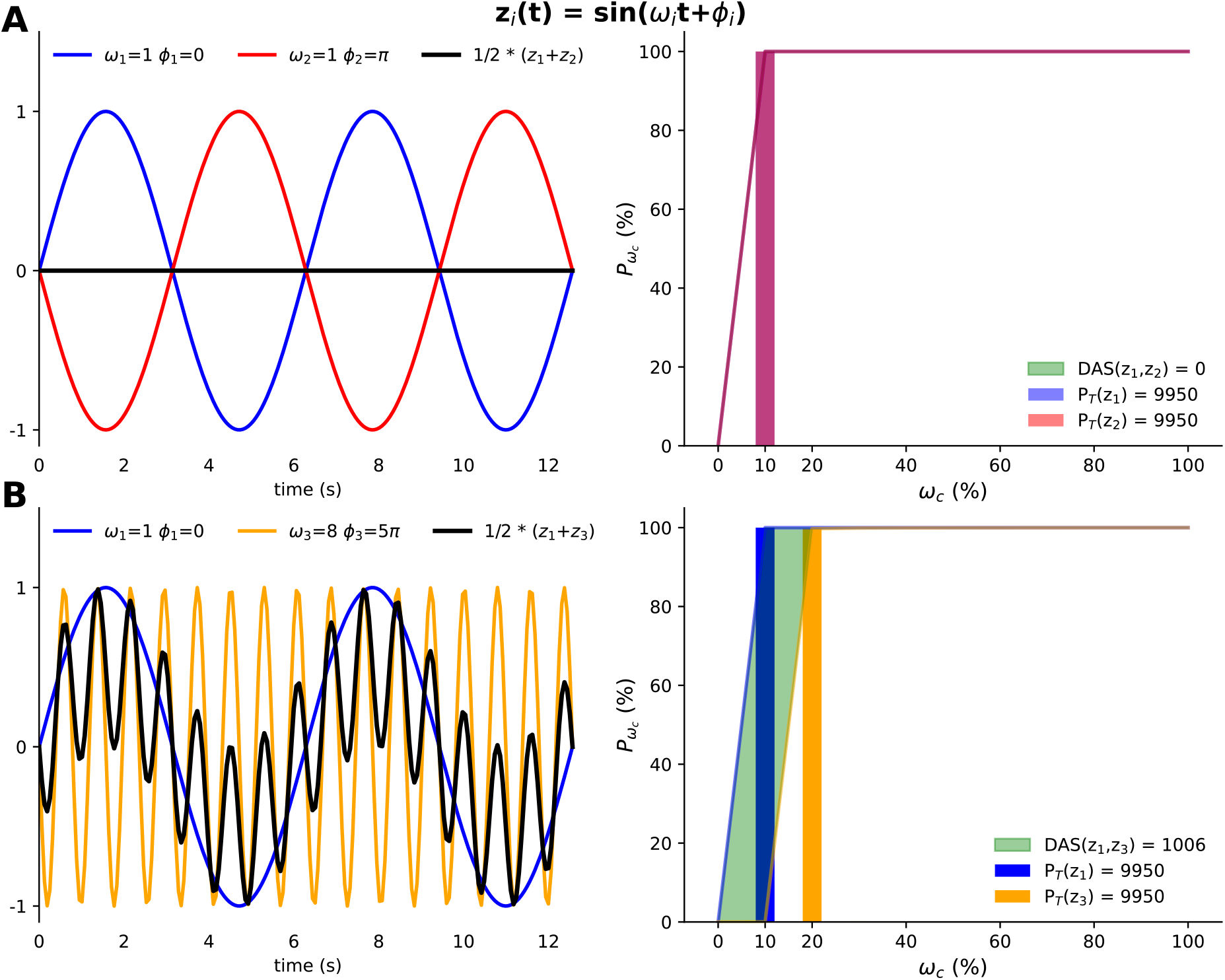
DAS example of 3 sinusoidal signals. **A)** LEFT Two sinusoidal signals with the same frequency and in complete dephasing. Their temporal correlation is exactly -1 and their mean is always zero and no coupled behavior can emerge. RIGHT The power spectrum, however, is identical (complete overlapping). Both signals, therefore, carry the same amount of power in the same frequency band. The lines represent the cumulative power distribution as defined in Eq. (3). In short, these two signals are indistinguishable since the dephasing might be a consequence of different starting time points. **B)** LEFT Two sinusoidal signals with different frequencies of oscillations and arbitrary dephasing can carry the same power. The mean of both signals is now the composition of both frequencies. RIGHT The spectrum of frequencies is, nonetheless, very different. Conventional distance metrics would only capture differences in the information content, but with the Dynamics Alteration Score (DAS) is able to distinguish which one of the signals displays faster or slower oscillations. The values of both the power and the DAS are entirely conditioned to the scale used (i.e., *DAS* ∈ [0,1] if we had not defined the distributions as percentages).

**Figure S2.**
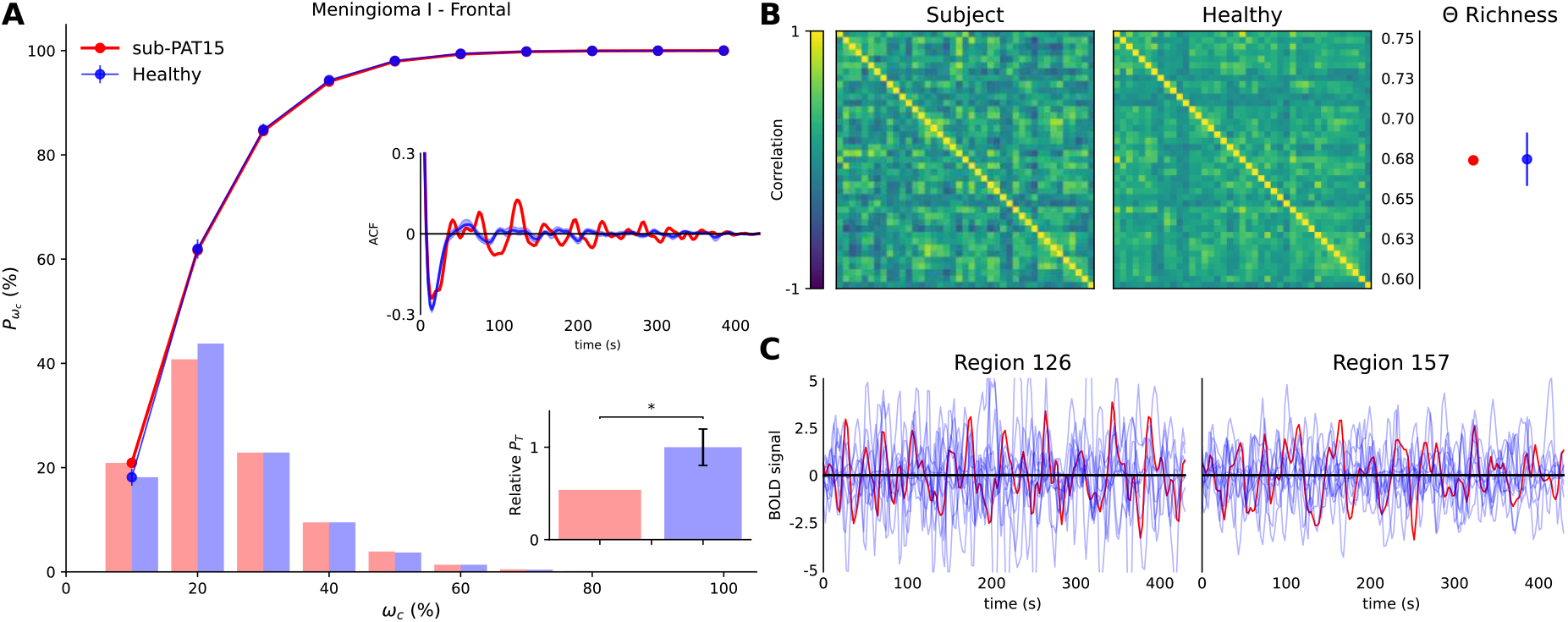
sub-PAT15 DMN. **A)** Cumulative power (lines), power distribution (bars), autocorrelation function (ACF) and total power relative to healthy subjects (histogram) for the patient (red) and mean of controls (blue) functional signals of the regions in the DMN. Error bars [mean ± SEM] indicate the results for the control population, while “*” code for statistical significance. **B)** LEFT Functional DMN from the same patient and mean of control subjects (see Methods). RIGHT Functional complexity as measured by the distribution of correlations for the patient’s network and the mean of control [mean ± SEM] (see Methods). **C)** Functional signal of two randomly selected regions from the same patient (red) as opposed to all the control subjects (light blue). No apparent difference is found between raw time series across regions, subjects and patients.

**Figure S3.**
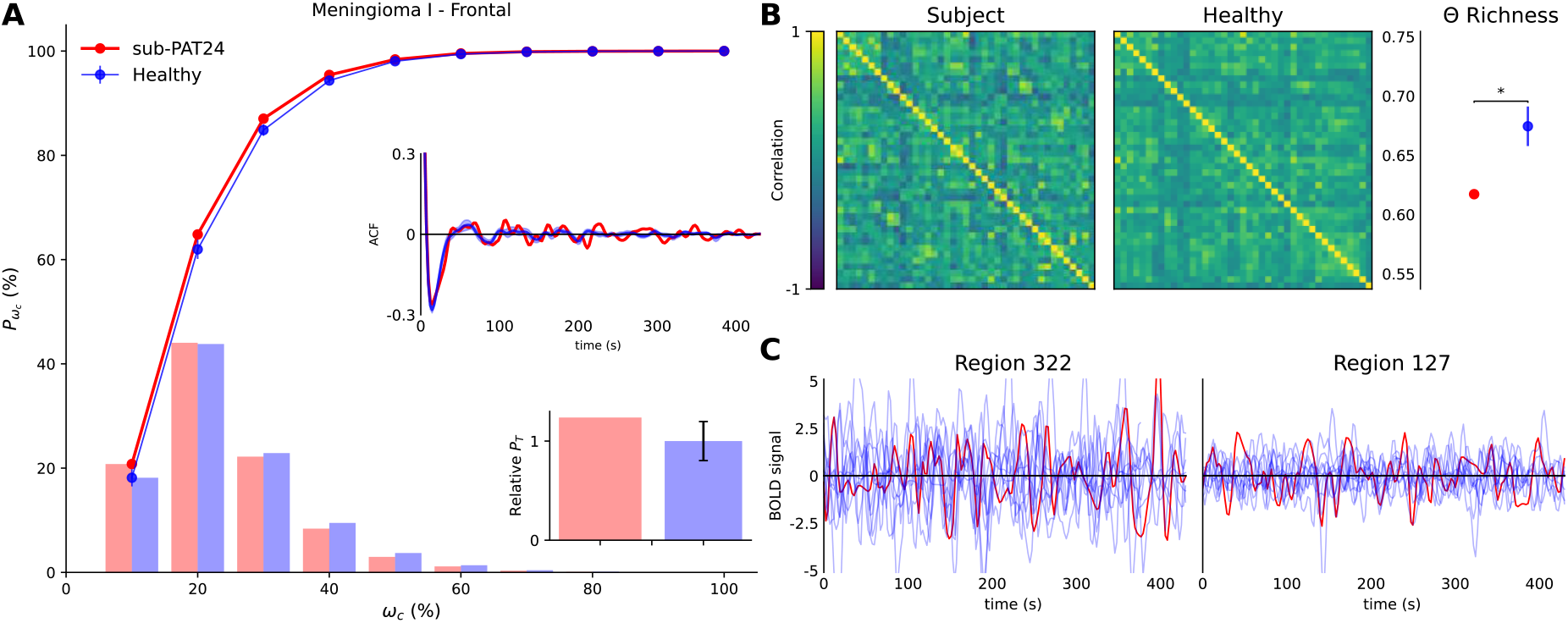
sub-PAT24 DMN. **A)** Cumulative power (lines), power distribution (bars), autocorrelation function (ACF) and total power relative to healthy subjects (histogram) for the patient (red) and mean of controls (blue) functional signals of the regions in the DMN. Error bars [mean ± SEM] indicate the results for the control population, while “*” code for statistical significance. **B)** LEFT Functional DMN from the same patient and mean of control subjects (see Methods). RIGHT Functional complexity as measured by the distribution of correlations for the patient’s network and the mean of control [mean ± SEM] (see Methods). **C)** Functional signal of two randomly selected regions from the same patient (red) as opposed to all the control subjects (light blue). No apparent difference is found between raw time series across regions, subjects and patients.

**Figure S4.**
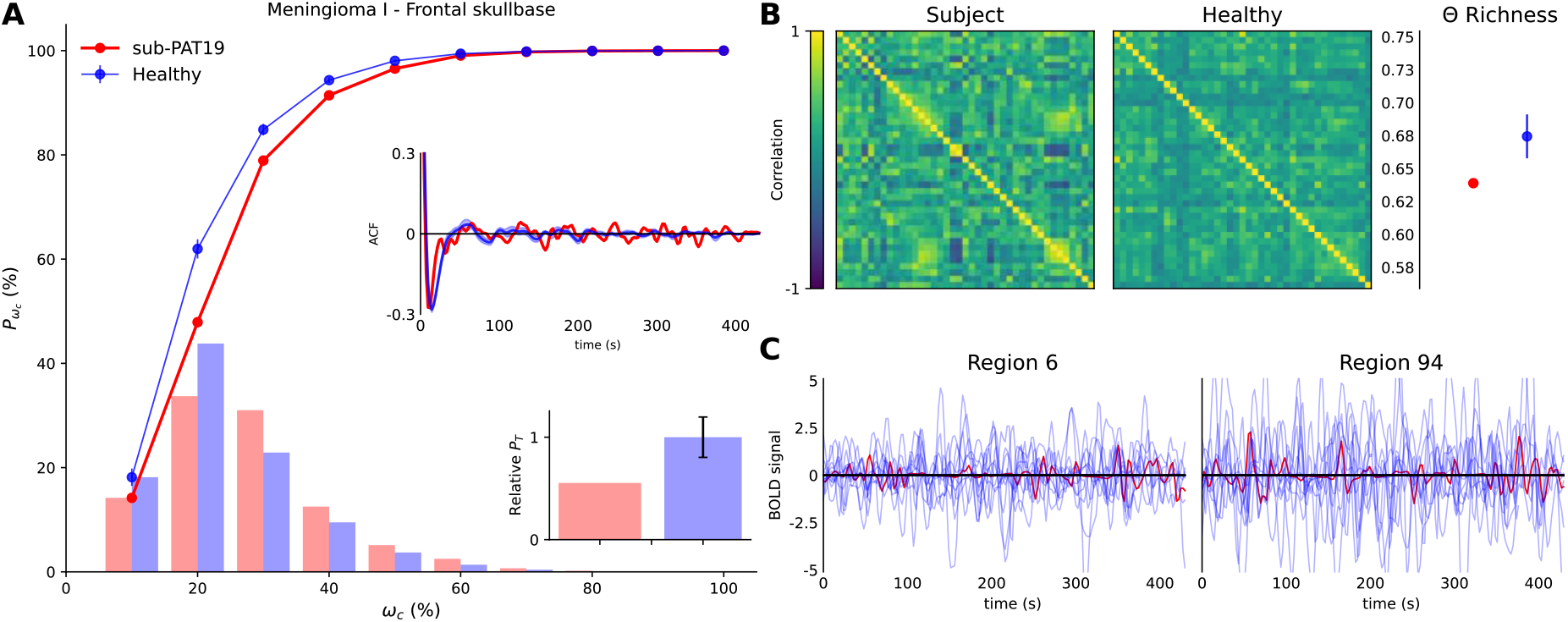
sub-PAT19 DMN. **A)** Cumulative power (lines), power distribution (bars), autocorrelation function (ACF) and total power relative to healthy subjects (histogram) for the patient (red) and mean of controls (blue) functional signals of the regions in the DMN. Error bars [mean ± SEM] indicate the results for the control population, while “*” code for statistical significance. **B)** LEFT Functional DMN from the same patient and mean of control subjects (see Methods). RIGHT Functional complexity as measured by the distribution of correlations for the patient’s network and the mean of control [mean ± SEM] (see Methods). **C)** Functional signal of two randomly selected regions from the same patient (red) as opposed to all the control subjects (light blue). No apparent difference is found between raw time series across regions, subjects and patients.

**Figure S5.**
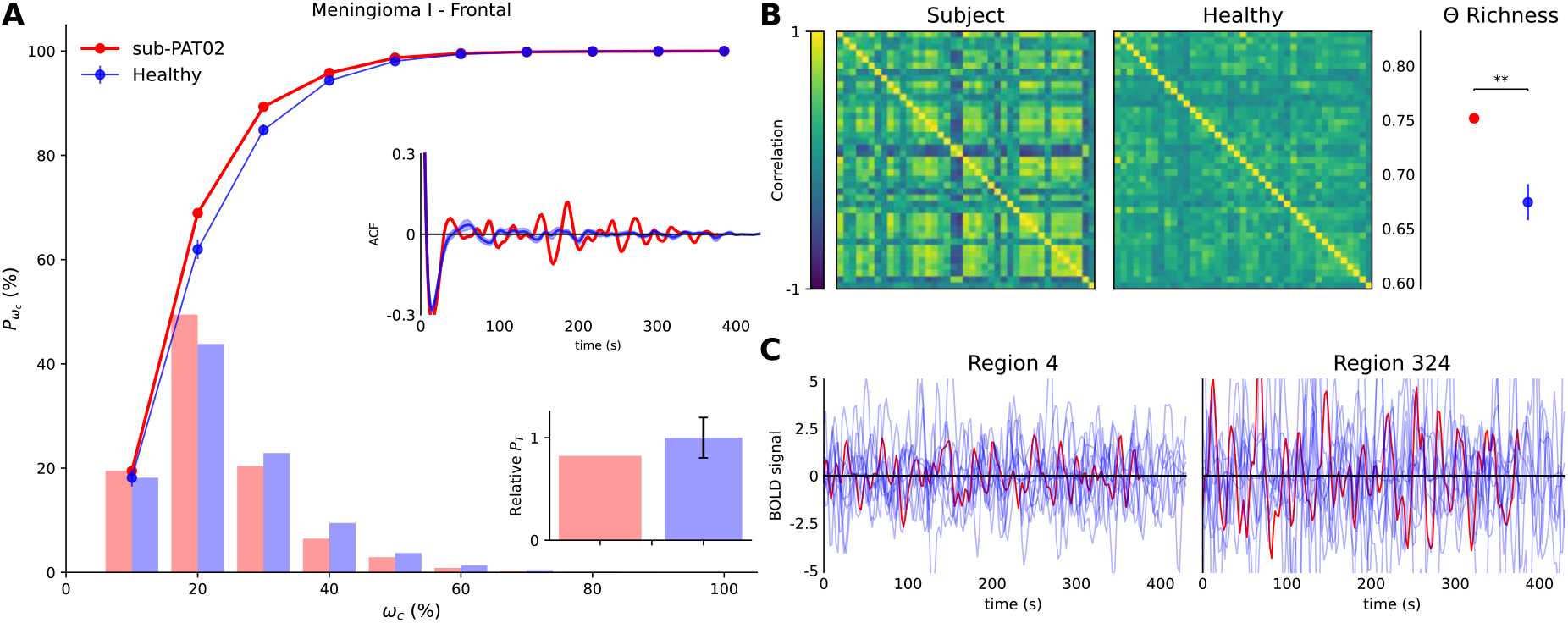
sub-PAT02 DMN. **A)** Cumulative power (lines), power distribution (bars), autocorrelation function (ACF) and total power relative to healthy subjects (histogram) for the patient (red) and mean of controls (blue) functional signals of the regions in the DMN. Error bars [mean ± SEM] indicate the results for the control population, while “*” code for statistical significance. **B)** LEFT Functional DMN from the same patient and mean of control subjects (see Methods). RIGHT Functional complexity as measured by the distribution of correlations for the patient’s network and the mean of control [mean ± SEM] (see Methods). **C)** Functional signal of two randomly selected regions from the same patient (red) as opposed to all the control subjects (light blue). No apparent difference is found between raw time series across regions, subjects and patients.

**Figure S6.**
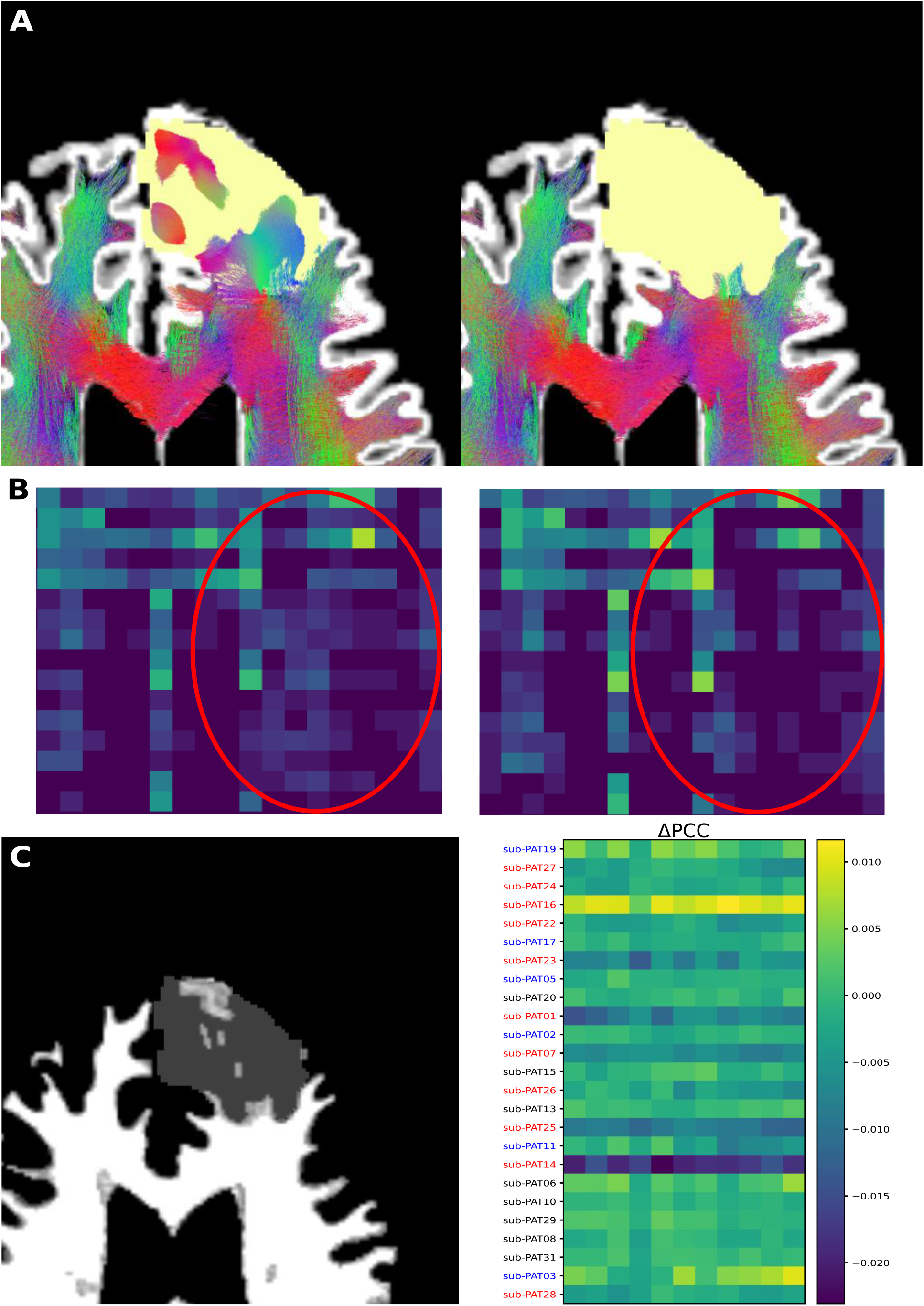
Combined Reconstruction Pipeline for sub-PAT29. **A)** LEFT Results from our combined pipeline and RIGHT results from a widely used multi-shell method. As expected, the reconstruction algorithm penetrates the tumor (Top row) creating connections that would not appear otherwise while maintaining those unaffected (Bottom row). **B)** LEFT White matter structure is heavily compromised in the presence of brain tumors (shaded region). RIGHT Differences in Pearson Correlation Coefficients between subjects and the 11 healthy controls. The combined approach increases similarity in most tumors but not in others. Colored subjects indicate a significant difference in PCC (red: p<0.01 and blue: p<0.05, U-test).

**Figure S7.**
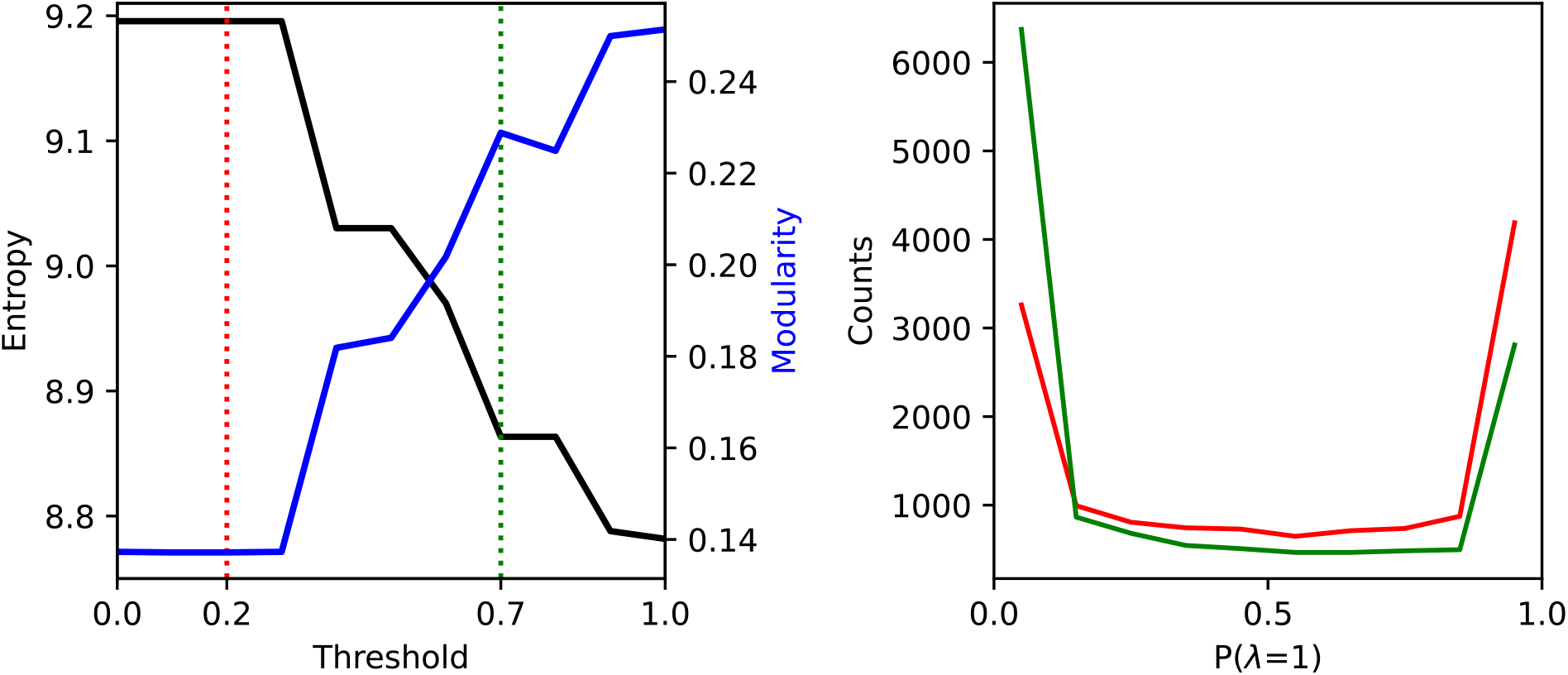
Choosing the threshold for the Anatomical Prior. LEFT Entropy and Modularity as a function of the selected threshold. These two properties where rather robust in certain intervals of the threshold value. Therefore, the choice of the threshold inside each one of these intervals was not critical to the performance of FCNET. Vertical dashed lines mark approximately the frontiers of the afore mentioned intervals. RIGHT Number of occurrences of the variable described by Eq. S1 for each one of the thresholds marked previously. Higher thresholds deleted a considerably large number f possible connections limiting the generalization power of FCNET.

## Footnotes

1 https://github.com/CyclotronResearchCentre/USwLesion

2 Note that the 3^rd^ version of the Automated Anatomically Labelled Atlas has 4 empty regions out 170 to maximize compatibility with the previous versions.

3 The repository will be updated soon. Do not hesitate to contact any of the two corresponding authors.

## Notes

### Competing Interest Statement

The authors have declared no competing interest.

### Summary of Updates

Fig S6 has been updated. References [31, 60] have been added in detriment of the previous [59, 82]. Abstract typo mistake has been fixed. Location of the figures within the manuscript has been made more clear.

